# SMYD5 is a ribosomal methyltransferase that catalyzes RPL40 lysine methylation to enhance translation output and promote hepatocellular carcinoma

**DOI:** 10.1101/2024.07.21.604450

**Authors:** Bisi Miao, Ling Ge, Chenxi He, Xinghao Wang, Jibo Wu, Xiang Li, Kun Chen, Jinkai Wan, Shenghui Xing, Lingnan Ren, Zhennan Shi, Natasha M. Flores, Zhihui Liang, Xinyi Xu, Ruoxin Wang, Jingdong Cheng, Yuanhui Mao, Pawel K. Mazur, Jiabin Cai, Fei Lan

## Abstract

While lysine methylation is well-known for regulating gene expression transcriptionally, its implications in translation have been largely uncharted. Trimethylation at lysine 22 (K22me3) on RPL40, a core ribosomal protein located in the GTPase activation center (GAC), was first reported 27 years ago. Yet, its methyltransferase and role in translation remain unexplored. Here, we report that SMYD5 has robust *in vitro* activity toward RPL40 K22 and primarily catalyzes RPL40 K22me3 in cells. The loss of SMYD5 and RPL40 K22me3 leads to reduced translation output and disturbed elongation as evidenced by increased ribosome collisions. SMYD5 and RPL40 K22me3 are upregulated in hepatocellular carcinoma (HCC) and negatively correlated with patient prognosis. Depleting SMYD5 renders HCC cells hypersensitivity to mTOR inhibition in both 2D and 3D cultures. Additionally, the loss of SMYD5 markedly inhibits HCC development and growth in both genetically engineered mouse (GEM) and patient-derived xenograft (PDX) models, with the inhibitory effect in the PDX model further enhanced by concurrent mTOR suppression. Our findings reveal a novel role of the SMYD5 and RPL40 K22me3 axis in translation elongation and highlight the therapeutic potential of targeting SMYD5 in HCC, particularly with concurrent mTOR inhibition. This work also broadens the understanding of lysine methylation, extending its significance from transcriptional regulation to translational control.

## Introduction

Protein lysine N^ε^-methylation plays a crucial role in various biological processes. While its impact on transcription regulation via histone proteins has been extensively studied over the past two decades or so, its role in translation remains largely unexplored. In this context, several mammalian ribosomal proteins, such as RPL4, RPL29, RPL40 and RPL36A, have been reported to contain lysine methylation^1–3^. Among these, RPL40 is a special ribosomal protein encoded by the *UBA52* gene. The precursor UBA52 protein is a fusion protein of 128 amino acids (aa), comprising an N-terminal fusion of a ubiquitin module (76 aa). After the removal of ubiquitin, the mature form of RPL40 is 52 aa in length and is one of the last components assembled into the 60S ribosomal subunit in cytoplasm^4,5^. In the mature 80S ribosome, RPL40 is located near the P stalk/GAC (<u>G</u>TPase <u>A</u>ctivation <u>C</u>enter) and SRL (Sarcin-Ricin loop), where elongation factors eEF1A and eEF2 bind. The elongation factors are crucial for recruiting peptidyl-tRNA to the A-site and for translocating it from A-site to P-site. RPL40 has been proposed to selectively regulate stress-related mRNA translation and confer resistance to elongation inhibitor Sordarin in yeasts^4,6^, suggesting an essential function of RPL40 in protein synthesis. Importantly, the trimethylation of K22 on RPL40 (RPL40 K22me3, equivalent to UBA52 K98me3) was identified by mass spectrometry analysis in rat liver 27 years ago^1^, and visualized in recent high-resolution ribosome structural studies^2,3^. However, the role of this modification in translation and ribosome function remains unclear.

SET and MYND domain-containing (SMYD) proteins constitute an evolutionarily conserved subfamily of lysine methyltransferases, characterized by a catalytic SET domain split by a MYND domain^7,8^. Among them, SMYD5 was recently reported to catalyze methylation of viral Tat protein and be involved in HIV infection^9^, as well as histone H3K36me3 at promoters and drive tumorigenesis in hepatocellular carcinoma (HCC)^10^, though the downstream mechanism is unclear. Indeed, data from the TCGA database indicate that SMYD5 mRNA levels are elevated in most cancer types with HCC being one of the most significant types (Supplementary information, Fig. S1a)^11,12^. Consistently, two recent multi-omics studies have found that both SMYD5 mRNA and protein levels are significantly elevated in HCC samples and are associated with poor clinical outcomes (Supplementary information, Fig. S1b-e)^13,14^.

In this study, we identify SMYD5 as a ribosomal lysine methyltransferase that predominantly catalyzes RPL40 K22me3. The SMYD5-RPL40 K22me3 axis is crucial for efficient translation elongation and overall protein synthesis. Deficiency of SMYD5 in HCC cancer cells leads to hypersensitivities to mTOR inhibitors, likely due to a compounded inhibitory effect on protein synthesis. Employing both *ex vivo* and *in vivo* HCC models, we further elucidate the critical role of SMYD5-mediated RPL40 K22me3 in sustaining cancer growth, especially under suppressed mTOR signaling. These findings underscore the potential of targeting SMYD5-RPL40 K22me3 axis as a therapeutic strategy for HCC patients.

## Results

### SMYD5 trimethylates ribosomal protein RPL40 *in vitro*

To search for RPL40 methyltransferase, we screened a panel of SET domain-containing lysine methyltransferases for activity against the purified recombinant RPL40 protein. Using *in vitro* methyltransferase (MTase) assay, we found that only SMYD5 methylated RPL40 (Fig. 1a). Using RPL40 peptide substrate (12-32 aa) and MALDI-TOF analyses, the Km and Kcat values were determined at 11.34 μM and 1,394 h^-1^ (Fig. 1b), with the value of Kcat/Km at ∼ 122.9 demonstrating a strong enzymatic activity compared to other known lysine methyltransferases, such as G9A^15^. We then confirmed that SMYD5 catalyzes trimethylation at the K22 site in both RPL40 recombinant protein and peptide (Fig. 1c; Supplementary information, Fig. S2a,b). The K22A replacement completely ablated the SMYD5 activity towards the peptide substrate (Fig. 1c). Further mutagenesis approaches using recombinant RPL40 protein as a substrate consistently demonstrated that SMYD5 methylation of RPL40 is specific to K22, as the K22A or K22R mutations ablated SMYD5 activity whereas K17R mutation did not (Fig. 1d). To exclude the possibility of co-purified contaminant associated with SMYD5 from E. coli, we confirmed that the predicted catalytically dead SMYD5 mutant (Y351A, based on homology to SMYD2, SMYD3 and G9A), could not methylate recombinant RPL40 (Fig. 1e). In contrast to previous reports suggesting that SMYD5 is a putative histone methyltransferase^10,16–19^, we have not detected any activity on mononucleosomes, histone H3 (1-21 aa, 22-44 aa) and H4 (10-30 aa) tail peptides (Fig. 1f; Supplementary information, Fig. S2c,d), while under the same condition, we could detect robust SMYD5 activity on RPL40 recombinant protein and peptide. Such results demonstrate that SMYD5 has strong *in vitro* methyltransferase activity toward RPL40 compared to histone substrates.

**Fig. 1:**
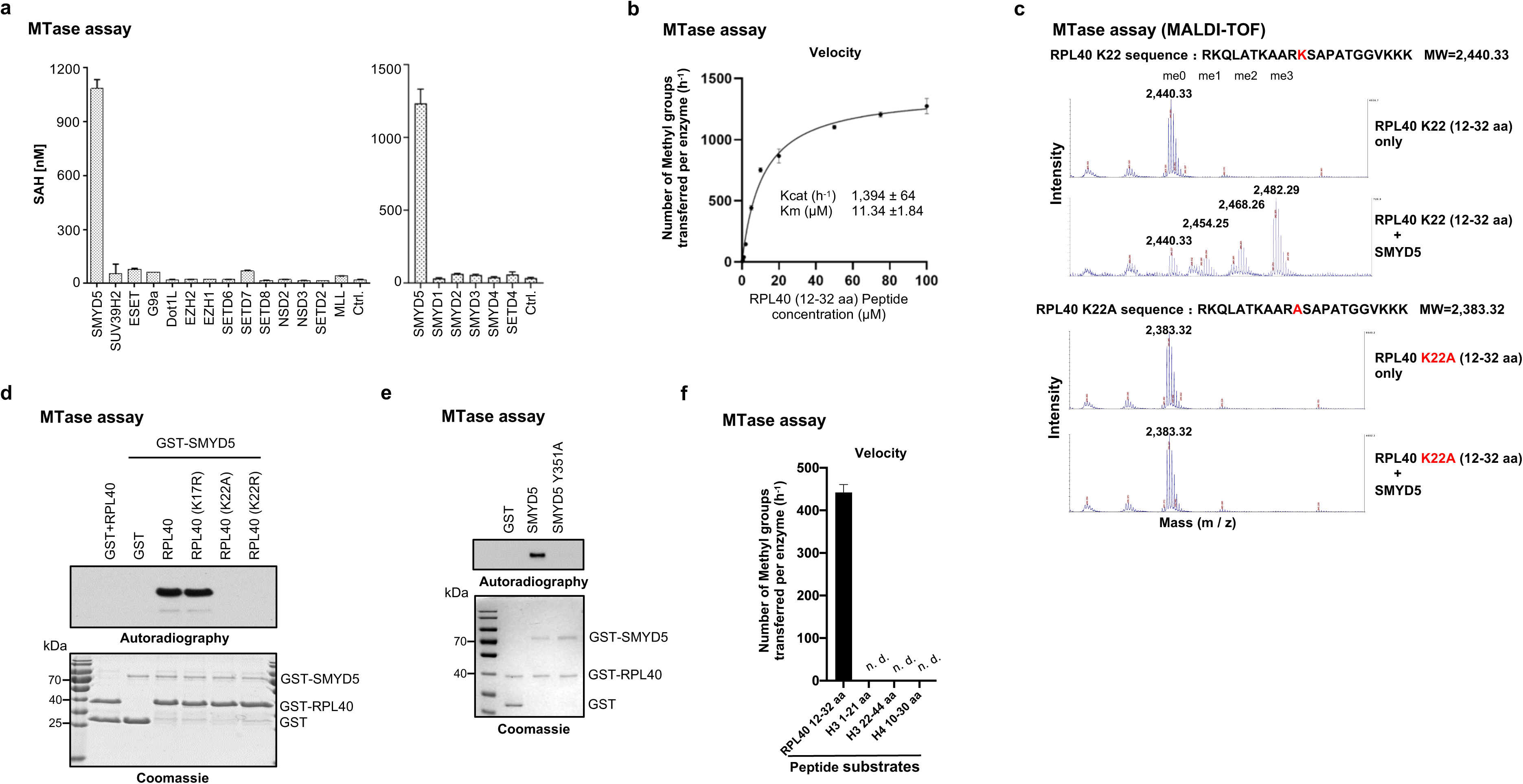
SMYD5 trimethylates ribosomal protein RPL40 *in vitro*. **a** *In vitro* methylation assays using recombinant RPL40 as substrate with a panel of indicated methyltransferases, with no enzyme as control (Ctrl.). The SAH converted by methyltransferase reactions from SAM was monitored by MTase-Glo Kit. All data are represented as mean ± SD from three biological replicates. Two-tailed unpaired t-test. **b** Measurement of the enzymatic parameters, Kcat and Km, using 20 nM purified recombinant SMYD5 protein and indicated concentrations of RPL40 peptide substrate (12-32 aa) for 30-minute reactions. The measurements were performed by MALDI-TOF. **c** MALDI-TOF analyses of the methyltransferase activities of the recombinant SMYD5 on RPL40 WT and K22A mutant peptides (12-32 aa). The concentrations of peptides in reactions were all at 5 μM and the concentration of recombinant SMYD5 protein was at 20 nM. **d** *In vitro* SMYD5 methylation assays with the indicated recombinant RPL40 wildtype and mutant proteins, and GST protein (as control). Top panel, autoradiogram of the methylation assays; bottom panel, coomassie blue staining of the protein components in the reactions. **e** *In vitro* methylation assays using recombinant RPL40 as substrate with the GST tagged wildtype and catalytic-dead (Y351A) SMYD5 proteins, and GST protein alone was used as a control. Top panel, autoradiogram of the methylation assays; bottom panel, Coomassie blue staining of the protein components use in the reactions. **f** *In vitro* measurements of SMYD5 methyltransferase activities with the indicated substrates. RPL40 peptide was compared to histone tail derived peptides. All peptides were at a concentration of 5 μM and reacted with a concentraion of 20 nM purified recombinant SMYD5 protein for 30 minutes. The measurements were performed by MALDI-TOF.

### RPL40 is the primary cytoplasmic substrate of SMYD5 in human cells

Congruous with our observation that SMYD5 lacks activity on histones, we observed that the endogenous and ectopically expressed SMYD5 predominantly localized to the cytoplasm in HeLa and Huh7 cells (Fig. 2a,b; Supplementary information, Fig. S3a,b). As RPL40 is known to be assembled to the 60S ribosomal subunit in cytoplasm^4,5^ and shows predominantly cytoplasmic localization (Supplementary information, Fig. S3c, image available from Human Protein Atlas), we next tested whether RPL40 is the major cytoplasmic substrate of SMYD5. To investigate this in an unbiased fashion, we carried out a methyltransferase (MTase) assay using total cytoplasmic protein lysates from either control or *SMYD5* knockout (KO) cells as substrates, and purified recombinant SMYD5 as enzyme with ^3^H-SAM (S-adenosyl methionine) as the methyl donor. Strikingly, we found a single strong autoradiographic signal that was specifically associated with the addition of SMYD5, using the lysate from SMYD5 KO cells (Fig. 2c). The signal migrated at molecular weight (MW) around and below 10 kDa, the predicted size of RPL40. Interestingly, we did not observe such a signal using cytoplasmic lysate from control HeLa cells under the same condition (Fig. 2c), indicating that the putative substrate might be close to be fully methylated in the control HeLa cells. Similar results were also observed in other human cell lines and mouse liver tissue (Supplementary information, Fig. S3d). We next performed MS/MS (tandem mass spectrometry) analyses of the gel-extracted HeLa proteins at ∼10 kDa and extensively analyzed the data to search for, 1) peptide sequences with potential lysine methylation and 2) with protein MW around 10 kDa. In this exercise, we filtered MS-derived peptides from 277 proteins, identifying only 6 hits that contained potential lysine methylation modifications. Notably, these hits included four ribosomal proteins, one of which was RPL40. These 6 peptides were synthesized and tested as substrates in an *in vitro* methylation assay with recombinant SMYD5 and ^3^H-SAM, and only the RPL40 peptide could be methylated (Supplementary information, Fig. S3e,f). Consistently, *in vitro* methylation assay using HeLa cytoplasmic lysate from cells depleted of RPL40 by RNAi as SMYD5 substrate resulted in a significant decrease in the ∼10 kDa autoradiographic signal when incubated with wild type but not the catalytically deficient recombinant SMYD5 (Fig. 2d). Altogether, these data indicate that RPL40 is the primary cytoplasmic substrate of SMYD5.

**Fig. 2:**
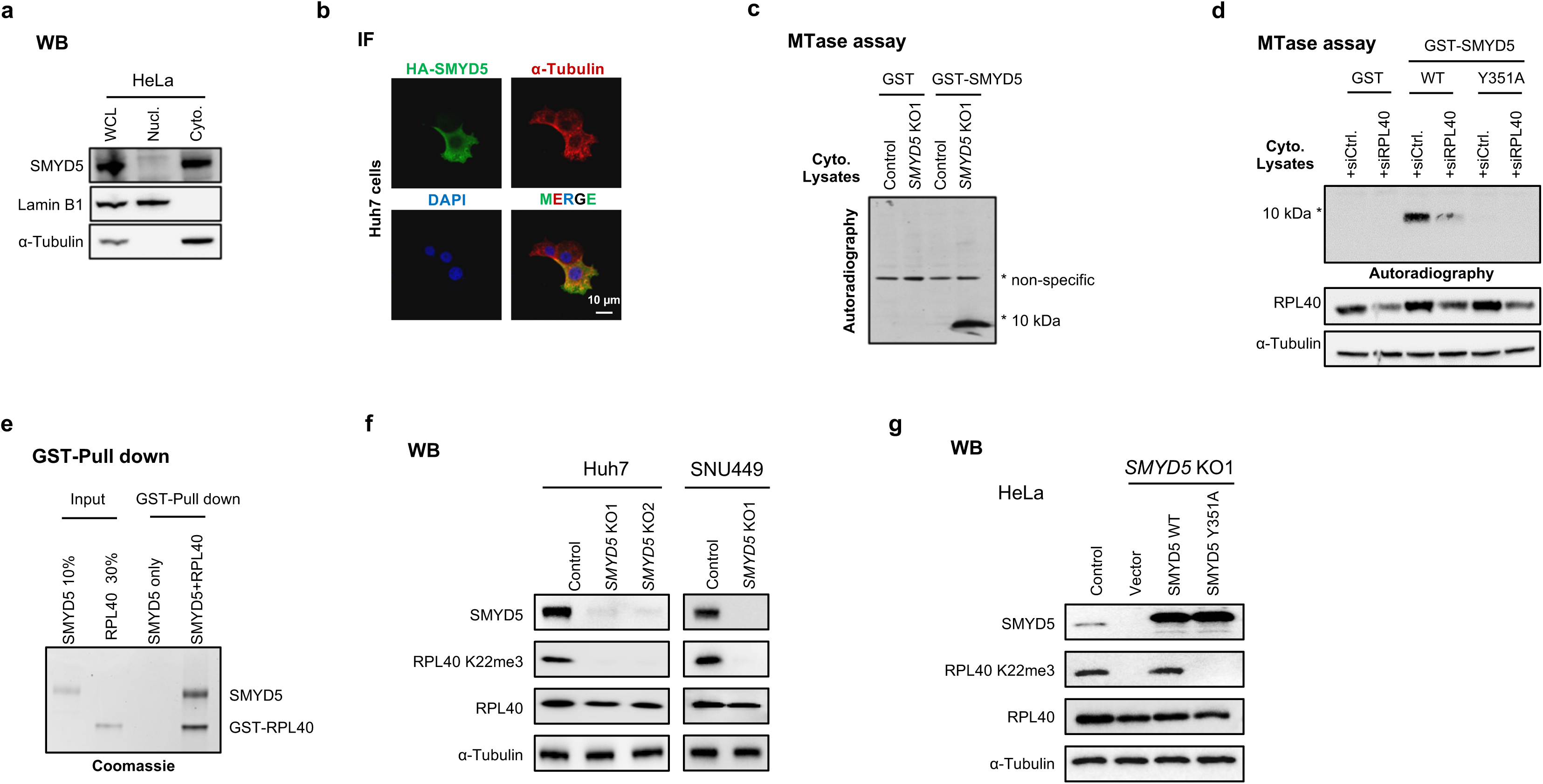
RPL40 is the primary cytoplasmic substrate of SMYD5 in human cells. **a** WB analyses of SMYD5 in the whole-cell lysate (WCL), cytoplasmic (C), and nuclear (N) fractions from HeLa cells. Lamin B1 and α-Tubulin were used as nuclear and cytoplasmic markers, respectively. **b** IF analyses of HA (green), α-Tubulin (red) and DAPI (blue) in Huh7 cells stably carrying HA-SMYD5 expression construct. **c** *In vitro* methylation reactions using recombinant GST-SMYD5 or GST (as a control) and cytoplasmic protein lysates from the control and *SMYD5* KO1 HeLa cells as substrates. * denoted the candidate substrate signal. **d** *In vitro* methylation reactions with recombinant GST tagged wildtype (WT), catalytic-dead SMYD5 (Y351A), or GST as control. Cytoplasmic protein lysates from control and RPL40 KD HeLa cells were used as substrates. Top panel, autoradiogram of the methylation assays; middle and bottom panel, WB analyses using indicated antibodies. **e** *In vitro* pull-down assays using recombinant GST tagged RPL40 and Flag tagged recombinant SMYD5. **f** WB analyses of SMYD5, RPL40 K22me3 and RPL40 in the indicated cell lines. *SMYD5* KO1 and KO2 cell lines of Huh7 and *SMYD5* KO1 cell line of SNU449 were generated by corresponding gRNAs in Methods. **g** WB analyses of SMYD5, RPL40 and RPL40 K22me3 in cells. *SMYD5* wildtype HeLa cells were used as the control. *SMYD5* KO1 HeLa cells were rescued by overexpressing empty vector, SMYD5 wildtype and SMYD5 mutant (Y351A) constructs.

To further consolidate the finding, we performed Flag-SMYD5 immunoprecipitation (IP) followed by mass spectrometry (MS) identification of SMYD5-interacting proteins. We found that the SMYD5 interactome contains many ribosome- and translation-related proteins (Supplementary information, Fig. S3g), with RPL40 scored as the second top binder. Although ribosomal proteins and translation factors are generally abundant and therefore often considered as contaminants in IP-MS experiments, the scores of unique peptides and peptide-spectrum match scores (PSMs) were of high confidence (greater than 55). The direct interaction between SMYD5 and RPL40 was also confirmed by GST pull-down assay (Fig. 2e).

To demonstrate whether SMYD5 is responsible for generating RPL40 K22me3 in cells, we raised a specific antibody against RPL40 K22me3 and validated it by dot blot assay (Supplementary information, Fig. S3h). Further validation was conducted using a HEK293T CRIPSR knock-in (KI) cell line in which lysine 22 of endogenous RPL40 was mutated to arginine (K22R), resulting in no detectable signal in RPL40 K22me3 immunoblotting (Supplementary information, Fig. S3i). Subsequently, we generated *SMYD5* knockout (KO) cells using CRISPR-Cas9 in HeLa, Huh7, SNU449 and HepG2 cell lines. We found that the RPL40 K22me3 signal was fully dependent on SMYD5 in all tested cell lines (Fig. 2f; Supplementary information, Fig. S3j). Furthermore, ectopical expression of wild type but not the Y351A mutant SMYD5, restored RPL40 K22me3 in *SMYD5* KO HeLa cells (Fig. 2g). We next found that the ratios of K22me3 / RPL40 were largely constant across multiple human cell lines and mouse tissues (Supplementary information, Fig. S3k,l). Targeted MS analyses found that RPL40 K22me3 was the major form in Huh7 cells and mouse liver (Supplementary information, Fig. S3m), consistent with previous report in rat liver^1^.

Together, these results support our *in vitro* findings and argue that SMYD5 is the physiologic enzyme that generates endogenous RPL40 K22me3, and the K22 methylated RPL40 is the major cellular form of RPL40 in the samples tested in this study.

### RPL40 K22me3 structural proximity to 28S rRNA in ribosomal GAC

To understand how SMYD5-RPL40 K22me3 may affect ribosome function, we analyzed the recently reported high-resolution Cryo-EM structure of the human mature ribosome^3^. As mentioned in introduction, RPL40 is located near the P stalk/GAC and SRL, where eEF1A and eEF2 bind (Supplementary information, Fig. S4a). The N-terminal alpha-helix (3-16 aa) associates with RPL9, which stabilizes the SRL, while the C-term domain, including K22, is inserted into a deep pocket formed by 28S rRNA helices H42, H89, H91 and H97, which are located next to the P stalk (Supplementary information, Fig. S4b middle). This part of RPL40 adopted a zinc finger structure with C20, C23, C34 and C39 chelating a zinc molecule (Supplementary information, Fig. S4b right). In the reported human ribosome structure (PDB 8GLP), the methyl electron density could readily be observed^3^ (Supplementary information, Fig. S4b right). The tri-methylated K22 (epsilon N) is positioned in a cleft between H42 and H89 of the 28S rRNA, with the closest distance to bases C4412 of H42 and G1945 of H89 at approximately 3.6 Å and 3.6 Å (Supplementary information, Fig. S4b right), respectively, a distance where van der Waals force may exist. Although the K22 side chain does not interact with translation factors, the H42 and H89 helices are known to interact with eEF2, eRF1 and A-site tRNA. Thus, it is plausible that without the trimethylation, the interaction and distance between RPL40 and 28S rRNA would be altered. Since the P stalk is a highly dynamic region during elongation, RPL40 methylation status could influence the overall P stalk conformation, which may impact the binding and release of elongation factors eEF1A and eEF2, potentially affecting the efficiency of protein synthesis.

### SMYD5 and RPL40 K22me3 affect polysome profiles and promote global translation output

We next investigated the impact of SMYD5 and RPL40 K22me3 on ribosome function and translation. First, using polysome profiling (ribosome sucrose gradient profiles), three consistent patterns in *SMYD5* KO HeLa and Huh7 cells were observed compared to the control cells (Supplementary information, Fig. S4c,d): 1) 40S, 60S and 80S fractions are largely unaffected by SMYD5 deletion; 2) *SMYD5* KO cells showed less heavy polysomes (> 5 ribosomes); 3) half-mer formation was observed in the light polysome and 80S fractions in the *SMYD5* KO cells, indicating altered active ribosome dynamics (also see discussion), similar to what has been reported in yeast with one copy deletion of the two RPL40 genes^4^. These results indicated that loss of SMYD5 affects active ribosome patterning, which may lead to reduced translation activity.

We then evaluated the impact of SMYD5 and RPL40 K22me3 on translation output by measuring newly synthesized proteins in Huh7 and SNU449 HCC cell lines using two independent methodologies: AHA click chemistry and Puromycin labeling (SUnSET). Both methods consistently demonstrated a 20-30% reduction in global translation output following SMYD5 deletion (Fig. 3a,b). A similar decrease in translation output was observed in the HEK293T K22R KI cell line, connecting the effect to the RPL40 K22 site (Fig. 3c). Further knockdown of SMYD5 in HEK293T K22R KI cells did not exacerbate this defect, supporting the idea that SMYD5’s role in translation primarily functions through RPL40 K22me3 modification (Fig. 3c).

**Fig. 3:**
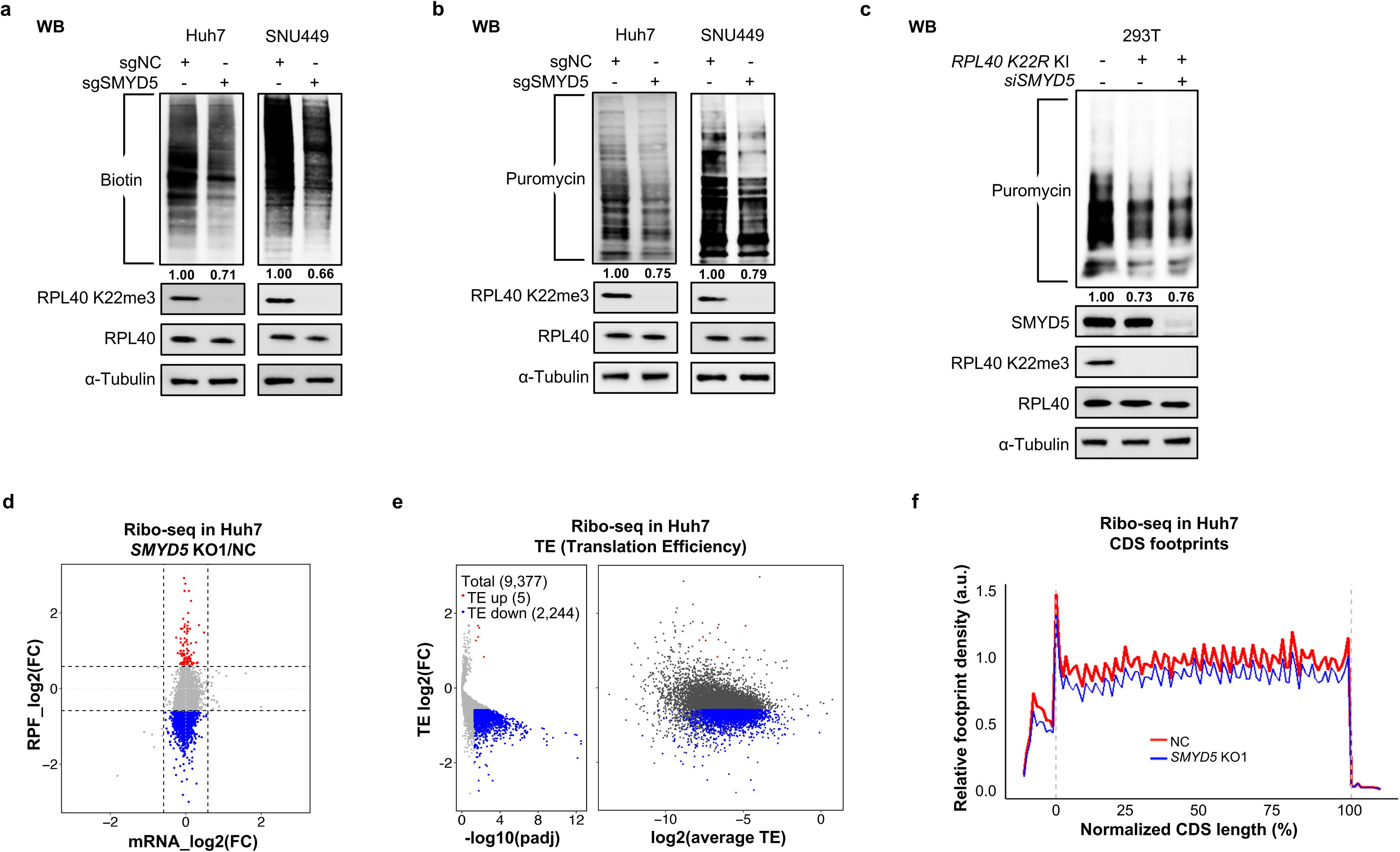
SMYD5 and RPL40 K22me3 promote global translation output by enhancing mRNA translation efficiency. **a-b** WB analyses of newly synthesized proteins in the negative control (NC) and SMYD5 depleted (KO1) Huh7 and SNU449 cell lines by AHA-click labeling (**a**) and puromycin (**b**) labeling approaches. The intensity of NC cells was normalized as 1.00 as indicated under different treatment. **c** Measurement of newly synthesized proteins in the wildtype (WT) and *RPL40 K22R* KI 293T cell lines by SUnset/Puromycin labeling approach. **d** Scatterplots of ribo-seq depicting the changes in transcription-level (x axis) and translation-level (y axis). The red and blue dots represent genes with stable mRNA levels but increase or decrease in RPF levels. RPF, ribosome protected footprint. FC (fold change) cutoff is 1.5. **e** Volcanoplots representing the translation efficiency (TE) changes from ribo-seq. Genes with FC >1.5 or < 1/1.5 and padj (adjusted-*P* value) < 0.05 were colored. Average TE is beween NC and *SMYD5* KO1, allowing for an evaluation of whether genes are highly/lowly translated. **f** Footprint of ribo-seq across normalized CDS reagions, normalized by mitochondrial RPF.

### SMYD5 enhances mRNA translation efficiency

To examine the SMYD5-denpendent translational landscape, we then conducted ribosome profiling sequencing (Ribo-seq) in Huh7 control and *SMYD5* KO cells^20^. A significant decrease in ribosome protected footprints (RPFs, i.e. ribosome protected fragments) was observed after *SMYD5* KO, indicating global reduced translation, while the transcriptome was largely unaffected (only few differentially expressed genes detected at fold change (FC) > 1.5) (Fig. 3d). We further calculated the translation efficiencies (TEs) of reliably detected mRNAs (9,377 transcripts), identifying a significant alteration in TEs among these transcripts (Fig. 3e; Supplementary information, Fig. S4e). Specifically, 300 mRNAs exhibited significantly decreased TE (FC > 2, padj (adjusted-p value) <0.05), and when the FC cutoff was adjusted to 1.5 fold, the number of mRNAs with decreased TE expanded to 2,244, while those with increased TE remained comparatively few (4 by 2-fold cutoff and 5 by 1.5-fold cutoff), indicating a broad impact of SMYD5 loss on the translatome. We also noticed that most mRNAs with significant TE reduction had a relatively higher initial TE level (Fig. 3e right). Additionally, ribosome footprint intensity tracks over coding DNA sequences (CDS) showed a global reduction in ribosome binding in *SMYD5* KO cells, without preferential changes in distribution (Fig. 3f). Moreover, no differential codon occupancy was detected between control and *SMYD5* KO cells (Supplementary information, Fig. S4f), suggesting that the translation reduction is globally uniform.

These results collectively confirm that the SMYD5-RPL40 K22me3 axis plays a crucial role in maintaining robust global translation output.

### Loss of SMYD5 - RPL40 K22me3 leads to elongation perturbation and hypersensitivity to translation inhibitors targeting A-site

As mentioned earlier, RPL40 is located near the GAC, a region where elongation factors bind and is closely related to A-site function (Supplementary information, Fig. S4a). The proximity leads us to hypothesize that the loss of SMYD5 and RPL40 K22me3 could disturb elongation. A common indicator of elongation perturbation is ribosome stalling and collision, frequently resulting in disome formation^21,22^. Consistent with our hypothesis, increased disome fractions were observed in Huh7 and SNU449 cell lines following SMYD5 depletion (Fig. 4a; Supplementary information, Fig. S5a). A similar effect was also observed in HEK293T *RPL40 K22R* KI cells (Supplementary information, Fig. S5b).

**Fig. 4:**
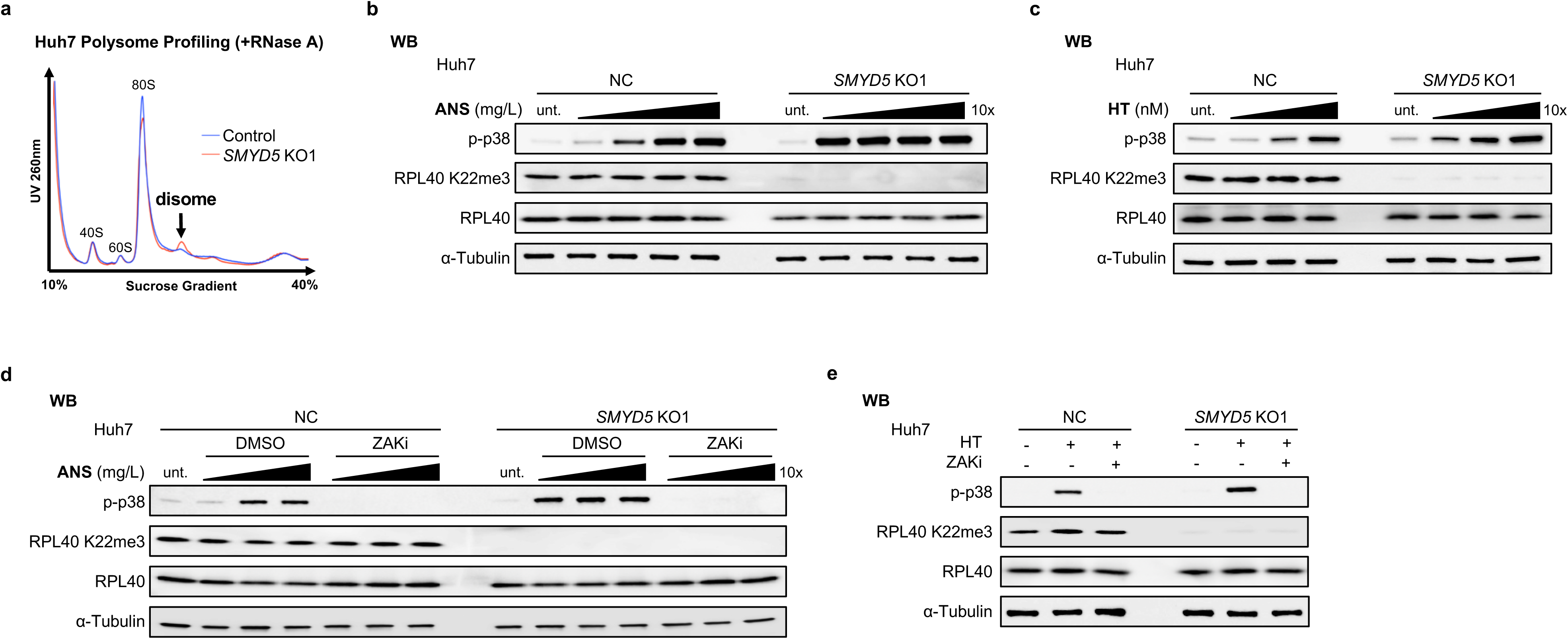
Loss of SMYD5 - RPL40 K22me3 leads to elongation perturbation and hypersensitivity to translation inhibitors targeting A-site. **a** Polysome profiles from lysates with RNase A treatment of NC and *SMYD5* KO1 Huh7 cell lines. Black arrows denoted the disomes. **b-c** WB analyses for phosphorylation of p38 in the NC and *SMYD5* KO1 Huh7 cell lines treated with Anisomycin (ANS) (0.001-1 mg/L, 15 min) or harringtonine (HT) (0.1-10 μM, 15 min). **d-e** WB analyses for phosphorylation of p38 in the NC and *SMYD5* KO1 Huh7 cell lines treated with ANS (0.01-1 mg/L, 15 min) or HT(10 μM, 15 min). Before ANS or HT treatment, the NC and *SMYD5* KO1 Huh7 cell lines were treated by DMSO or ZAK inhibitor M443 (5 μM, 1 hr) as indicated. Note: α-Tubulin was used as control for panels in all above and below WB analyses.

Recent studies have shown that significant ribosome stalling and collisions activate the ribosome-associated kinase ZAK, which in turn promotes p38 phosphorylation^22,23^, a key ribotoxic stress response (RSR) pathway. Although depletion of SMYD5 and RPL40 K22me3 alone was insufficient to activate p38, as indicated in Fig. 4b-e and Fig. S5c-f (compare the untreated condition), we speculated that the elongation perturbations caused by SMYD5 and RPL40 K22me3 loss might result in elevated sensitivities to RSR inducers. To test this idea, we measured the dosage-dependent p38 activation of several commonly used RSR inducers, including Anisomycin (ANS, targeting A-site), Harringtonine (HT, targeting A-site), Cycloheximide (CHX, targeting E-site), 254 nm UV irradiation (UVB) and Menadione (an inducer of <u>R</u>eactive <u>O</u>xygen <u>S</u>pecies, ROS) ^3,22–24^. We found that *SMYD5* KO Huh7 cells exhibited an ∼100-fold greater sensitivity to ANS and a ∼10-fold greater sensitivity to HT in activating p38 compared to control cells (Fig. 4b,c). Notably, this activation of p38 was entirely dependent on ZAK (Fig. 4d,e)^22,23^. Similarly, HEK293T K22R KI cells also demonstrated increased sensitivity to ANS treatments, directly linking the effect to the RPL40 K22 (Supplementary information, Fig. S5c).

Interestingly, SMYD5 loss in Huh7 cells did not significantly alter sensitivities to CHX, UVB and Menadione (Supplementary information, Fig. S5d-f), in contrast to the A-site targeting inhibitors, ANS and HT. These findings suggest that SMYD5-RPL40 K22me3 depletion specifically impairs ribosome fitness upon elongation perturbation and sensitizes elongating ribosomes to low levels of A-site targeting inhibitors. Such observed SMYD5-dependent elongation effects align with the physical positioning of RPL40 K22me3 near the GAC, which plays a crucial role in elongation and A-site function.

### SMYD5 loss sensitizes cancer cells to mTOR pathway blockade

Elevated protein synthesis and translation activity are hallmarks of cancer^25^, and as mentioned earlier, SMYD5 is frequently overexpressed in HCC and associated with poor clinical outcomes (Supplementary information, Fig. S1). We therefore postulated that SMYD5-mediated RPL40 K22me3 may promote HCC tumorigenesis. To explore this idea, we focused our investigation of SMYD5 function on three HCC cancer cell lines (Huh7, SNU449 and HepG2) and one model cell line HeLa.

Although SMYD5 depletion did not impact the proliferation of all four cell lines (Supplementary information, Fig. S6a) in normal cell culture conditions, we explored potential connections between SMYD5 and other pathways. To this end, we performed a comparative drug screen in control and *SMYD5* KO Huh7 cells using a library consisting of 172 small molecule inhibitors of major signaling, growth and epigenetics pathways (Fig. 5a; Supplementary information, Table S2). Interestingly, we found that the 4 top hits causing reduced fitness of *SMYD5* KO cells compared to the control cells, were inhibitors of the mTOR pathway, including Rapamycin, Torin1, Omipalisib (dual mTOR/PI3K inhibitor) and an apoptosis inducer Staurosporine that suppresses 4EBP1 phosphorylation (Fig. 5b). Such results raise the possibility that SMYD5 loss enhances mTOR dependency in Huh7 cells. Accordingly, viability assays showed elevated sensitivity of Huh7 cells to mTOR inhibitors, Torin1 and Rapamycin, upon SMYD5 loss, with IC50 values shifted roughly 5 times lower (Supplementary information, Fig. S6b). Consistently, rescue experiment demonstrated that K22R mutation was deficient in restoring Huh7 growth potential under Torin1 treatment as compared to the wild type RPL40 (Supplementary information, Fig. S6c). Elevated sensitivity to mTOR inhibition upon SMYD5 loss was also observed in SNU449, HepG2 and HeLa cells (Fig. 5c; Supplementary information, Fig. S6d), indicating a broader phenomenon.

**Fig. 5:**
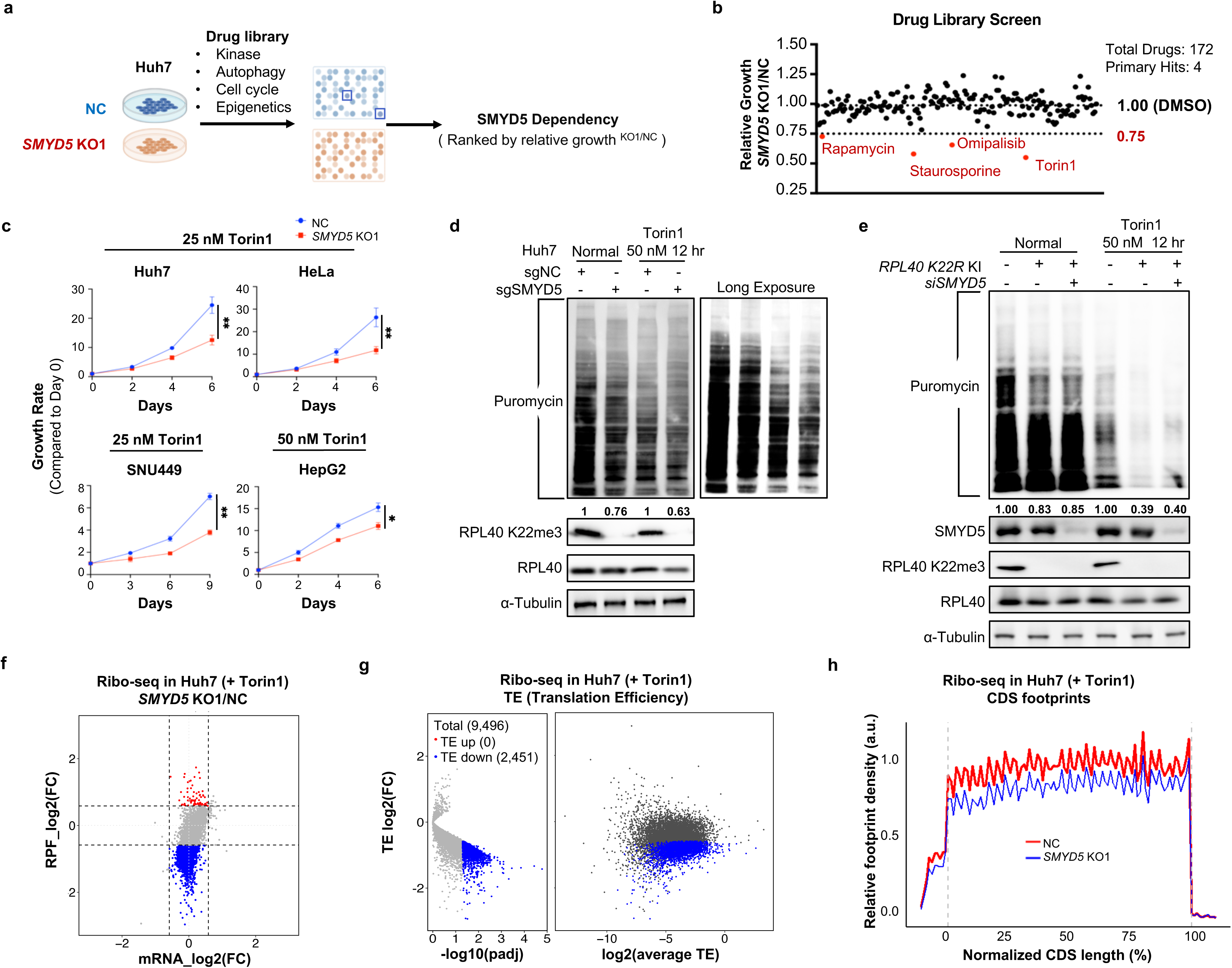
SMYD5 deletion sensitizes cancer cells to mTOR blockade. **a** Schematic of drugs screen in the control and *SMYD5* KO1 Huh7 cells. **b** Scatter plot of the fitness of *SMYD5* KO1 Huh7 cells compared to the control (NC) Huh7 cells under the treatment of individual drugs for 4 days. Data presented as relative growth rate of *SMYD5* KO1/NC cells. Four hits caused more than 25% growth retardation of *SMYD5* KO cells compared to the control cells were in red. Data were presented as the means from duplicated experiments. **c** Proliferation analyses of the control and SMYD5 depleted (KO1) Huh7, HeLa, SNU449 and HepG2 cells under the treatment of Torin1 at the indicated concentrations. Experiments were performed three times. **d** WB analyses of newly synthesized proteins in the NC and *SMYD5* KO1 Huh7 cell lines with or without Torin1 treatment by puromycin labeling approach. **e** WB analyses of newly synthesized proteins in the wildtype (WT), *RPL40 K22R* KI and *RPL40 K22R* KI with further knockdown (KD) of SMYD5 HEK293T cell lines with or without Torin1 treatment by puromycin labeling approach. **f** Scatterplots of ribo-seq depicting the changes between control and KO in transcription-level (x axis) and translation-level (y axis) with 50 nM Torin1 treatment for 12 hr. The red and blue dots represent genes with stable mRNA levels but increase or decrease in RPF levels. RPF, ribosome protected footprint. FC (fold change) cutoff is 1.5. **g** Volcanoplots of ribo-seq representing the translation efficiency (TE) changes. Genes with FC >1.5 or < 1/1.5 and padj (adjusted-*P* value) < 0.05 were colored. Average TE is beween NC and *SMYD5* KO1, allowing for an evaluation of whether genes are highly/lowly translated. **h** Footprint of ribo-seq across normalized CDS reagions under Torin1 treatment, normalized by mitochondrial RPF.

To assess the impact of SMYD5 loss on translation under mTOR suppression, we employed AHA click chemistry and SUnSET labeling techniques. Both methods confirmed a more pronounced SMYD5-dependent reduction in newly synthesized proteins in Huh7 and SNU449 cell lines under mTOR suppression compared to baseline conditions (Fig. 5d; Supplementary information, Fig. S6e-g; compare the ratios of reduction, lane4 / lane 3 vs lane 2 / lane 1). Consistently, enhanced decrease in newly synthesized proteins was observed in 293T *RPL40 K22R* KI cell lines under Torin1 treatment (Fig. 5e, compare the ratios of reduction, lane 5 / lane 4 vs lane 2 / lane 1). Further knockdown of SMYD5 in *RPL40 K22R* KI cells did not enlarge this defect, connecting the SMYD5-dependent effect to RPL40 K22 (Fig. 5e, compare the ratios of reduction, lane 6 / lane 4 vs lane 5 / lane 4). Additionally, ribo-seq analysis in Huh7 cells identified a significant number (491 out of a reliably detected total of 9,496, FC >2) of mRNAs with reduced TEs upon SMYD5 loss under Torin1 treatment, while no mRNAs showed significantly increased TEs. When the cutoff was lowered to FC 1.5, 2,451 mRNAs showed reduced TEs while still no mRNAs showed significantly increased TE (Fig. 5f,g; Supplementary information, Fig. S6h). In addition, similarly as shown in Fig. 3f, a greater global reduction of ribosome binding was observed over the CDS regions under Torin1 treatment in *SMYD5* KO cells (Fig. 5h). Again, no differential codon occupancy was detected under Torin1 treatment (Supplementary information, Fig. S6i), same as the untreated normal condition (Supplementary information, Fig. S4f).

Gene set enrichment analysis (GSEA) of ribo-seq reads revealed that G2M checkpoint, MYC targets, E2F targets and ribosome biogenesis were enriched in control cells compared to *SMYD5* KO upon Torin1 treatment (Supplementary information, Fig. S6j-l), in line with the growth defect of *SMYD5* KO cells under this condition (Fig. 5c). Consistent with these, under 12 hours of Torin1 treatment, both Huh7 and HeLa *SMYD5* KO cells showed less 40S, 60S, 80S and heavy polysome abundance compared to the control cells (Supplementary information, Fig. S6m). Western blot analyses also showed a moderate reduction in a few core ribosomal proteins in the *SMYD5* KO Huh7 cells upon mTOR suppression (Supplementary information, Fig. S6n), suggesting that the combined effect of SMYD5 loss and mTOR suppression might influence ribosome biogenesis, partly explaining the reduced growth potential of SMYD5-depleted cells under mTOR suppression. Taken together, these findings demonstrate that *SMYD5* KO cells experience substantial translation suppression when treated with Torin1.

### Loss of SMYD5 and mTOR inhibition synergize to suppress HCC development

To further explore SMYD5 function in HCC tumorigenesis when mTOR is suppressed, we monitored Huh7 cell proliferation using 3D soft agar and xenograft assays. We performed soft agar assay under 5% physioxia oxygen condition^26^ and found that the *SMYD5* KO Huh7 cells showed moderate defects in growth compared to the control cells, and the difference in growth potential was further pronounced upon mTOR inhibition (Fig. 6a). Similar results were observed *in vivo*, where 25 mg/kg Torin1 treatment led to significantly reduced xenograft growth of Huh7 *SMYD5* KO cells compared to the control Huh7 cells (Fig. 6b), while no statistical difference in xenograft growth was observed without mTOR inhibition (Fig. 6b). The Huh7 xenograft results were further supported by experiments using the human SNU449 cell line, which was established from poorly differentiated HCC. However, different from Huh7, SMYD5 ablation alone could readily attenuate xenograft growth of SNU449, and the growth was restored by complementation with wild type but not with catalytically inactive SMYD5 (Fig. 6c; Supplementary information, Fig. S7a). Importantly, the SMYD5 dependency of SNU499 xenograft tumor growth was further magnified by Torin1 treatment as well (Fig. 6c).

**Fig. 6:**
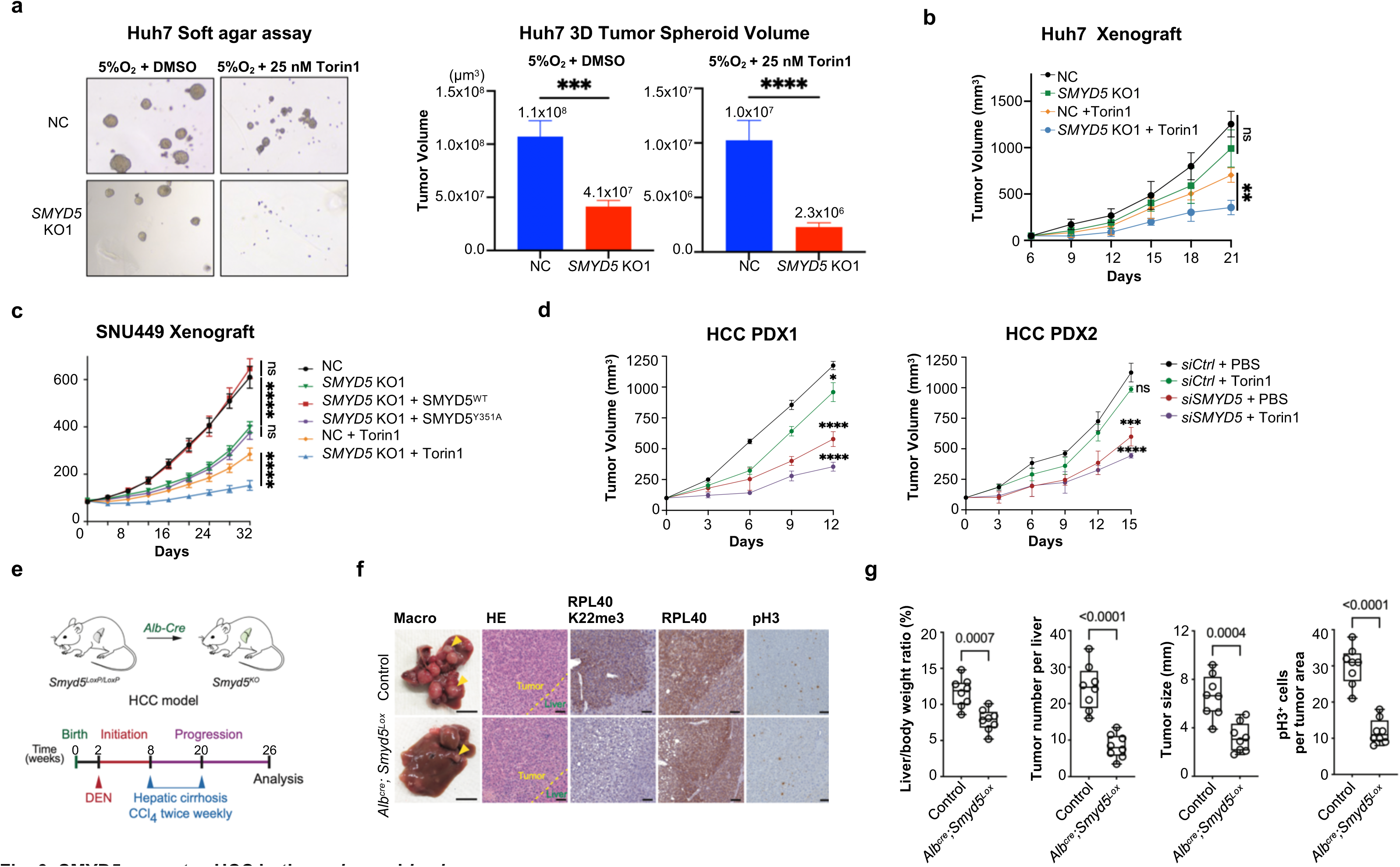
SMYD5 promotes HCC both *ex vivo* and *in vivo*. **a** Left, anchorage independent growth analyses using 3D soft agar assay of the control and *SMYD5* KO1 Huh7 cells under indicated treatments; Middle and right, quantification of the 3D spheroid volumes of the results was shown in the right panel. **b** Tumor volume quantification for Huh7 xenografts in nude mice (n= 4 for each group). The control and *SMYD5* KO1 Huh7 xenografts were both treated with PBS or Torin1 25 mg/kg daily. Data were represented as mean ± SEM. **c** Tumor volume quantification for SNU499 xenografts in NSG mice (n = 5 for each group). The control and *SMYD5* KO1 SNU449 cells, and the *SMYD5* KO1 SNU449 cells with overexpressing wildtype (WT) or catalytically deficient (mut) SMYD5 were used. 25 mg/kg Torin1 was used for intraperitoneal injection daily for the indicated groups. Data were represented as mean ± SEM. **d** Tumor volume quantification of HCC patients derived xenopgraft (PDX) in NSG mice (n = 4 for each group) from different two HCC patients. The siRNA that dissolved in the PBS with DMSO or 500nM Torin1 was used for intratumoral injection per three days. Data were represented as mean ± SEM. **e** Schematic illustrating animal HCC model utilized in the generation of liver-specific Smyd5 deletion; Lower, experimental design to assess effects of SMYD5 ablation on development of HCC with advanced liver fibrosis. **f** Left two columns, representative gross images of liver pathology (arrows indicated tumor nodules), HE-stained sections and immunohistochemical staining with indicated antibodies of tumors from control and SMYD5 depleted mice at 6 months of age (representative of n = 8 mice for each group), scale bars: 5 mm (whole mount) and 100 µm (histology); the rest columns, IHC representatives of RPL40 K22me3, total RPL40 and proliferation marker pH3 in control and SMYD5 depleted tumor samples. **g** Quantifications of liver/body weight ratio, tumor number, tumor size and pH3 positive cells in control and SMYD5 depleted tumor samples used in Fig. 6f. Boxes: 25th to 75th percentile, whiskers: min. to max., center line: median; n = 8 mice for each group.

To establish clinical significance of SMYD5 and RPL40 K22me3, we analyzed 202 human HCC patient samples from a Zhongshan Hospital cohort (Shanghai), performing IHC examinations to quantify the abundance of SMYD5, RPL40 and RPL40 K22me3. The analysis revealed predominant cytoplasmic staining of SMYD5, which was significantly elevated in 56 % of tumor (T) samples and lower in only 6 % of the tumor (T) samples compared to adjacent paratumor (P) tissues (Supplementary information, Fig. S7b,c). The intensity of SMYD5 staining inversely correlated with overall survival and disease-free survival rates (Supplementary information, Fig. S7d), consistent with previous reports (Supplementary information, Fig. S1)^13,14^. Like SMYD5, RPL40 K22me3 is elevated in 53.4% and attenuated in 5.0% of the tumor (T) samples compared to adjacent paratumor (P) tissues (Supplementary information, Fig. S7c). RPL40 also exhibited a similar, though less pronounced, trend (Supplementary information, Fig. S7c), consistent with the notion that the expression of ribosomal proteins is generally elevated in cancer^27^. Further analysis revealed that RPL40 K22me3 not only correlated strongly with RPL40 level, but also with SMYD5 level in these human HCC samples (Supplementary information, Fig. S7e,f). To further demonstrate clinical relevance, we then tested the SMYD5 effect in two HCC patient-derived xenograft (PDX) models. Using specially modified siRNAs for *in vivo* delivery (see Method), we depleted SMYD5 and RPL40 K22me3 in HCC PDX tumors via intratumoral injections. The results demonstrated that treatment of *SMYD5* siRNA alone readily resulted in significantly suppressed HCC PDX growth, with an even greater effect achieved by combined treatment of Torin1 (Fig. 6d; Supplementary information, Fig. S7g). Altogether, these findings underscore the critical role of SMYD5 and RPL40 K22me3 in promoting HCC progression.

### SMYD5 promotes HCC development *in vivo*

Human hepatocellular carcinoma (HCC) develops most often as a complication of liver cirrhosis. To investigate the role of SMYD5 in HCC tumorigenesis from the cirrhosis stage, we interbred conditional *Smyd5^loxP/loxP^* mutant mice with hepatocyte-specific Cre-recombinase strain (*Alb^Cre^*), resulting in specific *Smyd5* gene deletion in the liver (Fig. 6e). Animals with liver-specific SMYD5 depletion (*Alb^Cre^; Smyd5^loxP/loxP^*) developed normally without any notable phenotype, aligning with data from International Mouse Phenotyping Consortium (IMPC, Supplementary information, Table S3) showing that full body *Smyd5* knockout mice are viable, fertile and exhibits only minor phenotypes^28,29^. We utilized a model based on a two-stage chemical application to initiate and promote hepatocellular tumors in association with advanced liver fibrosis. The HCC model was induced by a single injection of genotoxic N-nitrosodiethylamine (DEN) followed by repeated administration of the pro-fibrogenic agent carbon tetrachloride (CCl_4_) (Fig. 6e). The control animals (*Alb^Cre^*) developed advanced HCC at 6 months of age. In contrast, SMYD5 ablation led to a significant reduction of overall tumor sizes and tumor burden with fewer tumor nodules noted (Fig. 6f,g). The liver/body weight ratio, a common indicator of total HCC tumor burden, was also significantly lower in *Smyd5* KO animals (Fig. 6f). In addition, depletion of SMYD5 resulted in a complete loss of RPL40 K22me3 staining and attenuation of tumor cancer cell proliferation (pH3) (Fig. 6f,g), in support of our mechanistic study. These observations are also consistent with the postulated role of SMYD5 and RPL40 K22me3 in HCC tumorigenesis.

## Discussion

Despite RPL40 K22 methylation has been identified in rat liver in 1997^1^, its role in ribosome function and translation control has not been explored due to the unawareness of its enzyme. Here, we identify SMYD5 as a ribosome protein methyltransferase responsible for RPL40 K22me3, allowing subsequent assessment of the role of this modification in translation. We demonstrate that the SMYD5 methylation on RPL40 K22 enhances global translation output and is required for proper translation elongation. Loss of SMYD5 and methylation on RPL40 leads to elevated ribosome collisions and hypersensitivity to translation inhibitors targeting A-site (Fig. 4a-c; Supplementary information, Fig. S5a-c). RPL40 is positioned near GAC where elongation factors, eEF1A and eEF2, bind. Conformational dynamics at this region is foreseeable during elongation, which involves cycles of A-site tRNA loading and A-to P-site transition. The increased sensitivities particularly to translation inhibitors targeting A-site indicating that RPL40 K22me3 is important in maintaining proper GAC conformation and dynamics, likely through the interaction of the methylated sidechain of K22 with 28S rRNA (Supplementary information, Fig. S4a,b). Of note, our ribo-seq analyses find that SMYD5 loss causes strong TE reduction of thousands of translating mRNA, without involving codon-specific effects (Fig. 3d,e; Supplementary information, Fig. S4e,f). Based on these findings, we conclude that the role of the SMYD5-RPL40 K22me3 axis is required for general elongation but not selective translation. In addition, SMYD5 may potentially impact initiation and 60S assembly as indicated by observed half-mers (Supplementary information, Fig. S4c,d), which requires further investigation.

The observation of elevated ribosome collisions or stalling upon loss of SMYD5-RPL40 K22me3 is of interest (Fig. 4a; Supplementary information, Fig. S5a,b). Recently, ribosome collisions/stalling and subsequent activation of RSR pathway have been connected to a variety of upstream stresses and cellular processes, such as inflammation, ageing and liver metabolic disorder^24,30^. Of note, interfering a ribosome modifier/modification causing ribosome collisions has not been reported yet. Our findings suggest that SMYD5 and RPL40 K22me3 axis enhances ribosome fitness under conditions of low-level elongation stress, particularly associated with A-site perturbation. Whether the axis is an integral part of the ribosome stress sensing pathway is an intriguing question for future studies. As yeast ribosomes lack this modification (Supplementary information, Fig. S7h)^31^, and close SMYD5 homolog start to appear from insects. It is plausible that SMYD5 and RPL40 K22me3 have evolved in higher organisms for ribosomes to confront more complex environmental stresses. Through in-depth structural comparison using current available structures with enough high resolutions (Supplementary information, Fig. S8), we have not observed significant alteration of RPL40 and its K22 side chain due to translation factor binding. Therefore, the detailed mechanism of how RPL40 K22me3 affects A-site function and GAC/P-stalk dynamics remains unclear and our findings encourage future investigation toward this direction.

Elevated ribosome biogenesis and translation output are hallmarks of cancer^27^, and overexpression of ribosome proteins has been linked to enhanced cancer proliferation and metastasis^32,33^. Recent studies have expanded the scope to include rRNA modifications in cancer^34,35^, and some modifications exhibit variable levels between cancer and normal tissues, suggesting a mechanism for ribosome heterogeneity^36^. Yet, the role of ribosome lysine methylations and their modifiers in cancer remains underexplored. Although our data suggest that RPL40 is close to fully methylated in human cell lines and mouse tissues tested (Supplementary information, Fig. S3d,k-m), future studies should thoroughly investigate the physiological/pathological ranges of RPL40 K22me3 and the regulation of SMYD5 itself. In this context, the observed elevation of SMYD5, RPL40 and RPL40 K22me3 in HCC, correlated with poor prognosis, underscoring the clinical relevance of our discoveries. HCC often arises from complex metabolic, epigenomic, and proteomic reprogramming, rather than from direct genetic mutations^37^, making effective targeted therapies for HCC scarce. While mTOR inhibitors have shown limited efficacy in treating HCC^38,39^, our findings demonstrate that disrupting SMYD5-mediated methylation of RPL40 K22me3 enhances the sensitivity of HCC cells to mTOR inhibition. This indicates that a dual-targeting approach could surpass the effectiveness of mTOR inhibition alone.

Our SNU449 xenograft and patient-derived xenograft (PDX) HCC models demonstrate that SMYD5 depletion alone significantly suppresses cancer growth, and renders them hypersensitivity to mTOR suppression (Fig. 6b,c). Furthermore, the GEM model of cirrhosis-associated HCC with hepatocyte-specific SMYD5 knockout indicated that the SMYD5-RPL40 K22me3 pathway is critical in tumor initiation and progression (Fig. 6e-g). Compared to its limited effect in cell culture assays (Supplementary information, Fig. S6a), the observed robust phenotype of SMYD5 ablation in animal models could possibly be explained by the constrained mTOR activity in vivo, due to limited nutrition and oxygen availability^40^. Our study thus nominates SMYD5 as a novel therapeutic target for HCC, whose efficacy could be further enhanced in combination with mTOR suppression. Of note, given that siRNA approaches have recently been shown to be effective for liver-related gene targeting in humans, our findings from HCC PDX models are of significant clinical value (Fig. 6d; Supplementary information, Fig. S7g). Importantly, since SMYD5 is also elevated in other cancer types (TCGA, Supplementary information, Fig. S1a), future research should focus on developing specific inhibitors or modulators of SMYD5 as a general approach for cancer therapy.

Targeting the translational machinery -- a strategy that has proven successful in hematopoietic malignancies with the ribosome-targeting agent (Homoharringtonine, also called Omacetaxine) developed from Chinese medicine and granted accelerated approval by FDA in 2012^41–43^ -- remains underexplored with ribosomal modifiers. Here, we find that targeting SMYD5 may exhibit favorable safety profiles, as evidenced by the normal liver development and viability of *Smyd5* knockout mice (Supplementary information, Table S3). These suggest that targeting ribosome modifier SMYD5 might present fewer toxicity issues compared to direct targeting ribosomes.

In contrast to the well-established role of histone lysine methylation in transcription regulation, the function of lysine methylation in ribosomal proteins during translation remains largely unexplored. Similar to nucleosomes, ribosomes are ribonucleoprotein complexes made of positively charged small ribosome proteins, enriched with lysine and arginine, and negatively charged rRNAs. Such physicochemical properties lay the foundation for analogous epigenetic mechanisms to exist and to directly influence ribosomes. Indeed, in addition to RPL40, previous studies have identified lysine methylation on mammalian RPL4, RPL29 and RPL36A^1–3,44^. It is unclear whether lysine methylation occurs on more ribosomal proteins, particularly those located at flexible regulatory regions that current structure analyses cannot visualize. Our study thus calls for future research in this emerging area, aiming to expand the understanding of lysine methylation’s significance beyond transcription to encompass ribosome biology.

## Acknowledgements

This work is supported in part by grants from National Natural Science Foundation of China and the National Key Research and Development Program of China to F.L. (NSFC 31925010 and 2021YFA1300100), J.Cai (NSFC 82272703) and C.H. (NSFC 32300461).

## Competing Interests

Fei Lan is a scientific co-founder and stockholder of Active Motif Shanghai, Inc. and Alternative Bio, Inc., Pawel K. Mazur is a scientific co-founder, consultant and stockholder of Amplified Medicines, Inc. and Ikena Oncology, Inc. and a consultant and stockholder of Alternative bio, Inc.

## Author Contributions

B.M., L.G., C.H., X.W. and J.W. contributed equally to the study. They were responsible for the experimental design, conduction, data process and manuscript preparation. B.M., X.W., C.H. and K.C. identified enzymatic activity and completed most biochemistry assays and cellular characterization under the guidance from F.L. L.G., C.H. and S.X. conducted drug library screen and most cancer cell growth analyses. L.G., and C.H. performed polysome profiling and ribo-seq with the help of Y. M and J. Cheng. X. L. and C.H. analyzed ribo-seq data and J. Cheng analyzed the structure. J.Cai conducted patient cancer sample IHC analyses and P.K.W. is responsible for mouse work of SNU449 and HCC model. P.K.W., J.Cai and F.L. were equally responsible for project overseeing and planning, data interpretation and manuscript preparation.

---

**Supplementary information, Fig. S1.**
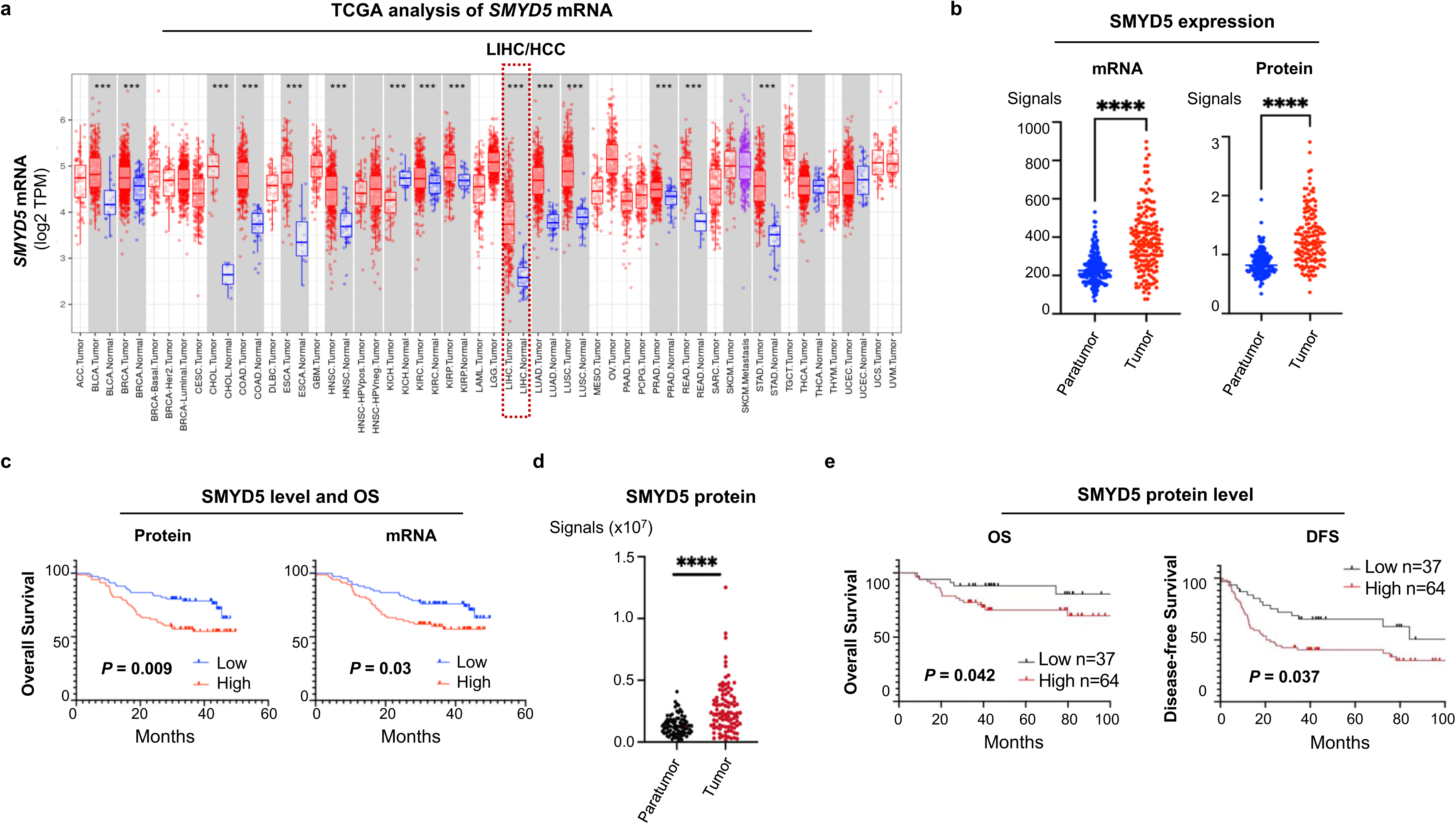
*SMYD5* mRNA and protein levels are elevated in multiple cancers including HCC. **a** TCGA database analyses showed elevated *SMYD5* mRNA levels in the cancer tissues (red) compared to the corresponding normal controls (blue), especially in hepatocellular carcinoma (HCC, in the red dashed box). **b** Strip plot analyses of *SMYD5* mRNA and protein levels in the tumor tissues of HCC samples compared to corresponding paratumor control. Data were retrieved from Qiang Gao, Cell, 2019. **c** Kaplan-Meier’s analyses of the overall survival (OS) of patients (n = 160) divided by SMYD5 high and low (mRNA and protein levels, respectively) in HCC. SMYD5 expression levels in HCC were negatively correlated with patients survival with indicated *P* values. Data were retrieved from Qiang Gao, Cell, 2019. **d** Strip plot analyses of SMYD5 protein levels in the tumor tissues of HCC samples compared to corresponding paratumor controls. Data were retrieved from Ying Jiang, Nature, 2019. **e** Kaplan-Meier’s analyses of the overall survival (OS) and disease-free survival (DFS) of patients divided by SMYD5 high and low (protein level) in HCC. SMYD5 expression levels in HCC were negatively correlated with OS and DFS with indicated *P* values. Data were retrieved from Ying Jiang, Nature, 2019. Note: All *P*-values above and below were determined by a two-tailed unpaired t-test and \**P < 0.05,* ** *P* < 0.01, *** *P* < 0.001, \*\*\*\**P* < 0.0001.

**Supplementary information, Fig. S2.**
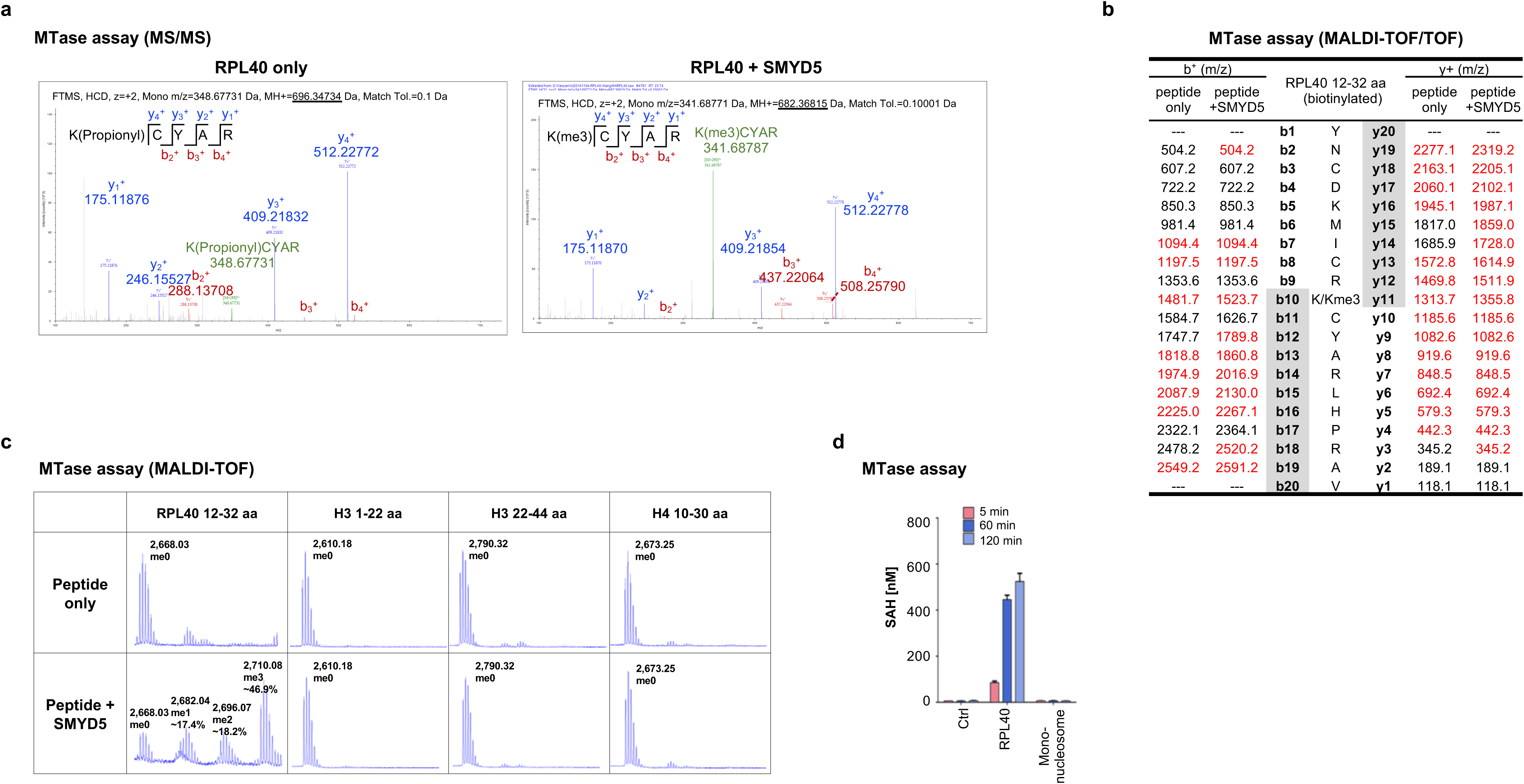
Related to Fig. 1. **a** MS/MS determination of the methylated lysine after propionylation, using recombinant full length RPL40 proteins as substrates. The detected b and y ions were indicated with the predicted molecular weights. **b** MALDI-TOF/TOF determination of the methylated lysine. The predicted b and y ions were listed, and the detected b and y ions were in red. **c** MALDI-TOF analyses of the methyltransferase activities of the recombinant SMYD5 with the indicated peptide substrates. Molecular weights were labelled and data quantifications were presented in Fig.1f. The reactions were performed for 1 hour and the concentrations of peptides were all at 5 μM and the concentration of recombinant SMYD5 protein was at 20 nM. **d** Recombinant RPL40 protein was compared to mononucleosome as substrates, and the SAH converted by methyltransferase reactions from SAM was monitored by MTase-Glo Kit at the indicated time points.

**Supplementary information, Fig. S3.**
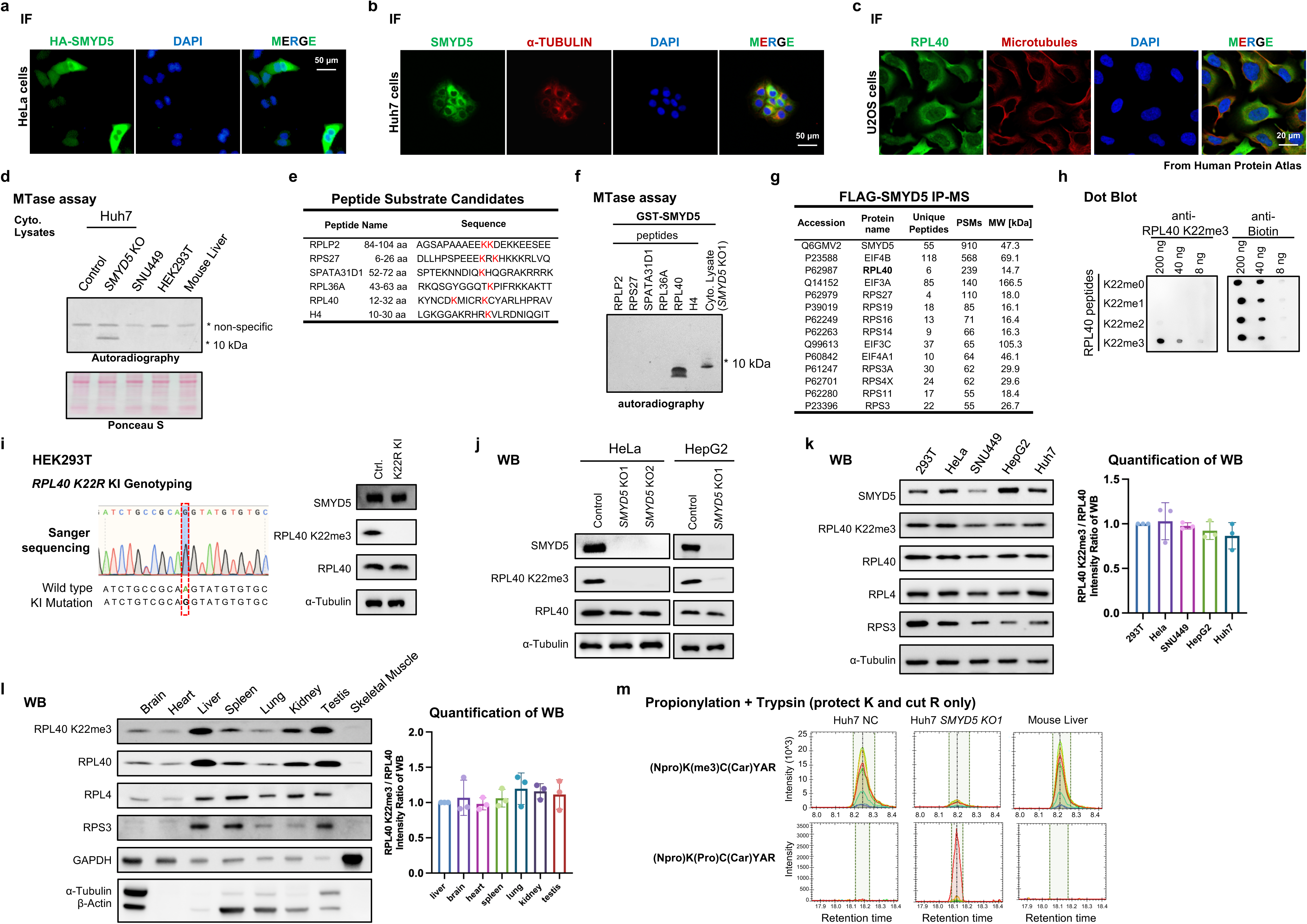
Related to Fig. 2. **a-b** IF analyses of exogenous HA tagged SMYD5 in HeLa cells and endogenous SMYD5 in Huh7 cells. **c** IF analyses of endogenous UBA52 in U2OS cells from Human Protein Atlas. The images are avaliable from v23.0.proteinatlas.org. URL: https://www.proteinatlas.org/ENSG00000221983UBA52/subcellular **d** In vitro HMT methylation reactions using recombinant GST-SMYD5 and lysates from human cell lines or mouse normal liver as substrates. **e** Potential methylated sequences and the candidate modified lysines were shown in red. **f** Autoradiography analyses of *in vitro* methylation reactions with recombinant SMYD5 using the indicated peptides as substrates. Cytoplasmic protein lysates from *SMYD5* KO1 cells were used as control (last lane). * denoted the candidate substrate signal from cytoplasmic lysate. **g** Top proteins identified by mass spectrometry (MS) in Flag-SMYD5 immunoprecipitation (IP), ranked by the values of peptide-spectrum match (PSM). **h** Specificity test of the RPL40 K22me3 antibody by dot blot using the indicated peptides. All peptides were modified by biotin and the anti-biotin signals were used as loading controls (right). **i** Left, genotyping of *RPL40 K22R* KI HEK293T cell line. Right, WB analyses of SMYD5, RPL40 K22me3 and RPL40 in the control and *RPL40 K22R* KI HEK293T cells. **j** WB analyses of SMYD5, RPL40 K22me3 and RPL40 in the indicated cell lines. *SMYD5* KO1 and KO2 cell lines of HeLa and *SMYD5* KO1 cell line of HepG2 were generated by corresponding gRNAs in Methods. **k-l** WB analyses of SMYD5, RPL40 K22me3, RPL40 and some core ribosome proteins in the indicated cell lines and mouse normal tissues. α-Tubulin, β-Actin or GAPDH were used as control. Intensities of WB were quantified in the right panels, relative ratios were normalized to 293T (**k**) or liver (**l**). **m** PRM ion transitions from methylated or unmethylated K22 precursor peptide by Propionylation + Trypsin strategy (see methods).

**Supplementary information, Fig. S4.**
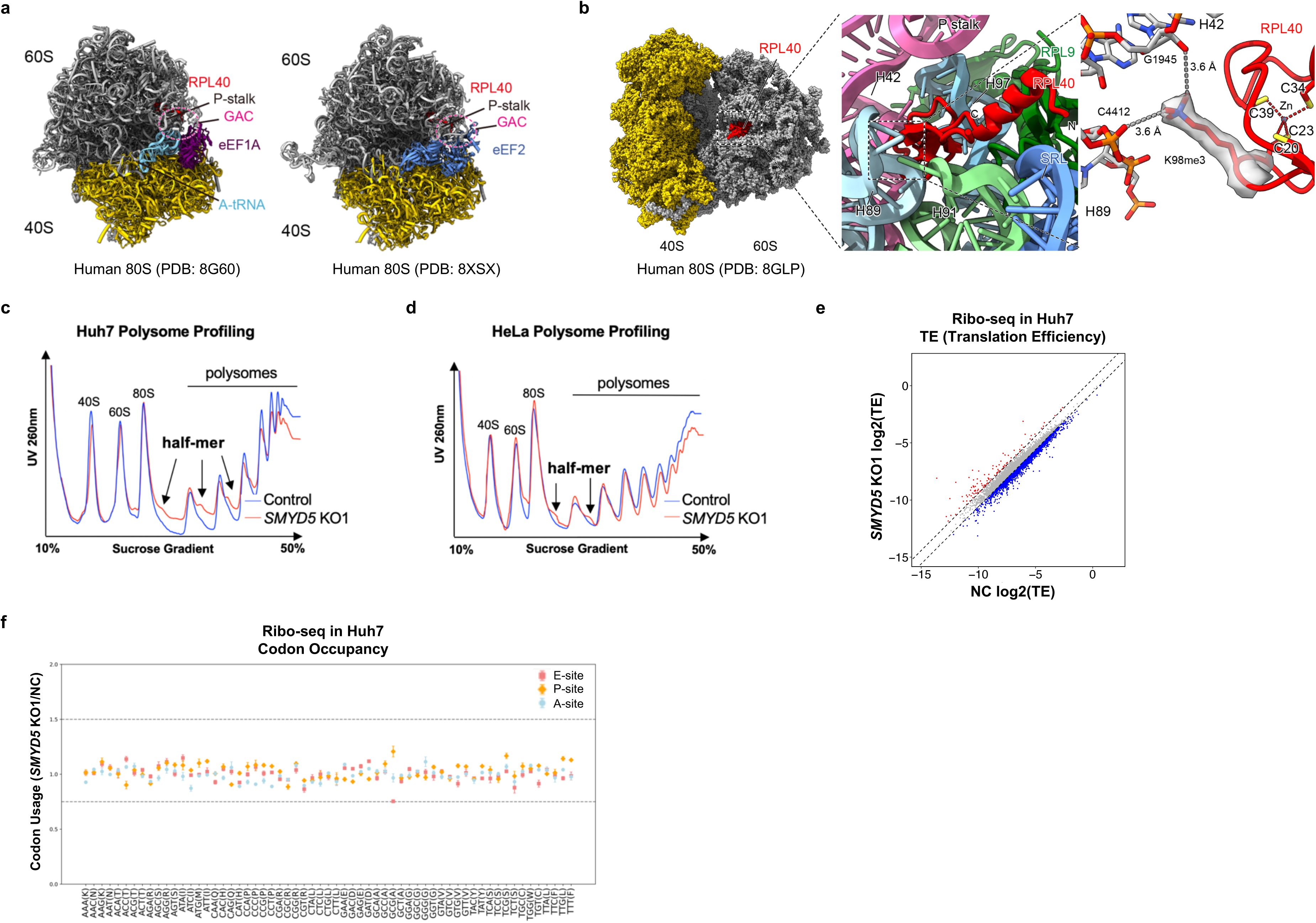
RPL40 K22me3 structural proximity to 28S rRNA in ribosomal GAC and affect polysome profiles. **a** Overview of RPL40 and its lysine 22 (K22) in a ribosome structure. Functional regions such as the P stalk, GAC, L1 stalk, PTC (peptidyl transferase center), CP (central protuberance) are indicated. Translation factors are highlighted in distinct colors. **b** Left, RPL40 in complex with human 80S ribosome; middle, a closer view of RPL40 in complex with 28S rRNA; right, distances between methylated K22 and C4412 of the H89 loop (28S rRNA) and methylated K22 and G1945 of the H42 loop (28S rRNA) are indicated. The analysis of the ribosome structure and its density map is based on PDB 8GLP. K22me3 is marked as “K98me3 of eL40” in the original PDB file. Different helix are highlighted in different colors. **c-d** Polysome profiles of the NC and *SMYD5* KO1 Huh7 (**c**) and HeLa (**d**) cell lines. Black arrows denoted the half-mers. **e** Scatterplots of ribo-seq showing translation efficiency (TE) correlation between NC and *SMYD5* KO1 Huh7 cells. Genes with fold change > 1.5 or < 1/1.5 were colored. **f** Codon bias analysis of ribo-seq between control and *SMYD5* KO1 Huh7 cells. The fold changes of each codon occupancy were calculated, and the dashed lines marked 1.5-fold threshold.

**Supplementary information, Fig. S5.**
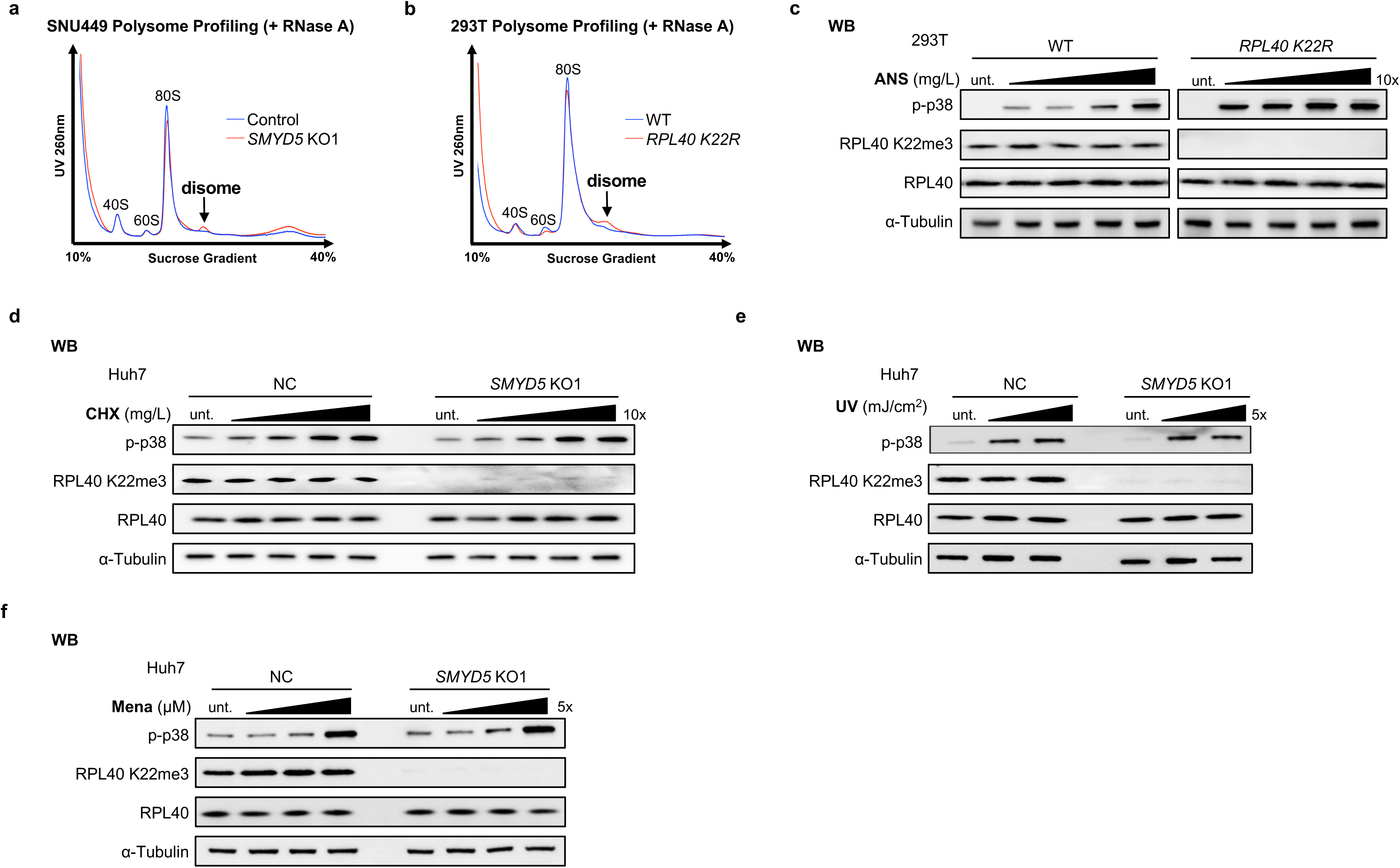
Related to Fig. 4. **a-b** Polysome profiles from lysates with RNase A treatment of NC and *SMYD5* KO1 SNU449 (**a**) and WT and *RPL40 K22R* KI 293T(**b**) cell lines. Black arrows denoted the disomes. **c** WB analyses for phosphorylation of p38 in the WT and *RPL40 K22R* KI 293T cell lines treated with ANS (0.01-10 mg/L, 15 min). **d-f** WB analyses for phosphorylation of p38 in the NC and *SMYD5* KO1 Huh7 cell lines treated with cycloheximide (CHX) (0.1-100 mg/L, 15 min) (**d**), 254 nm UV irradiation (1-5 mJ/cm^2, recovery 15 min) (**e**) or menadione (Mena) (1-25 μM, 15 min) (**f**) .

**Supplementary information, Fig. S6.**
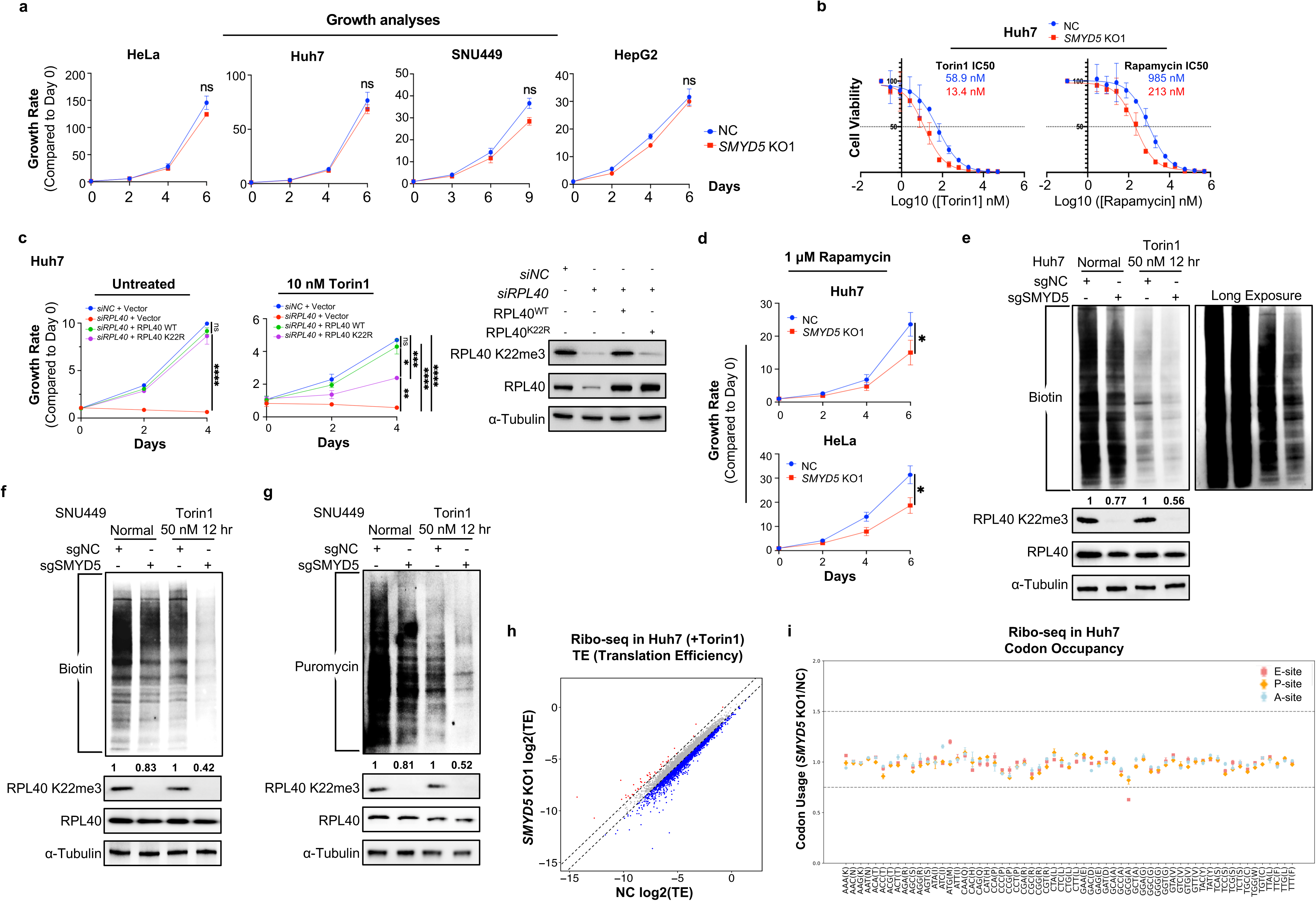
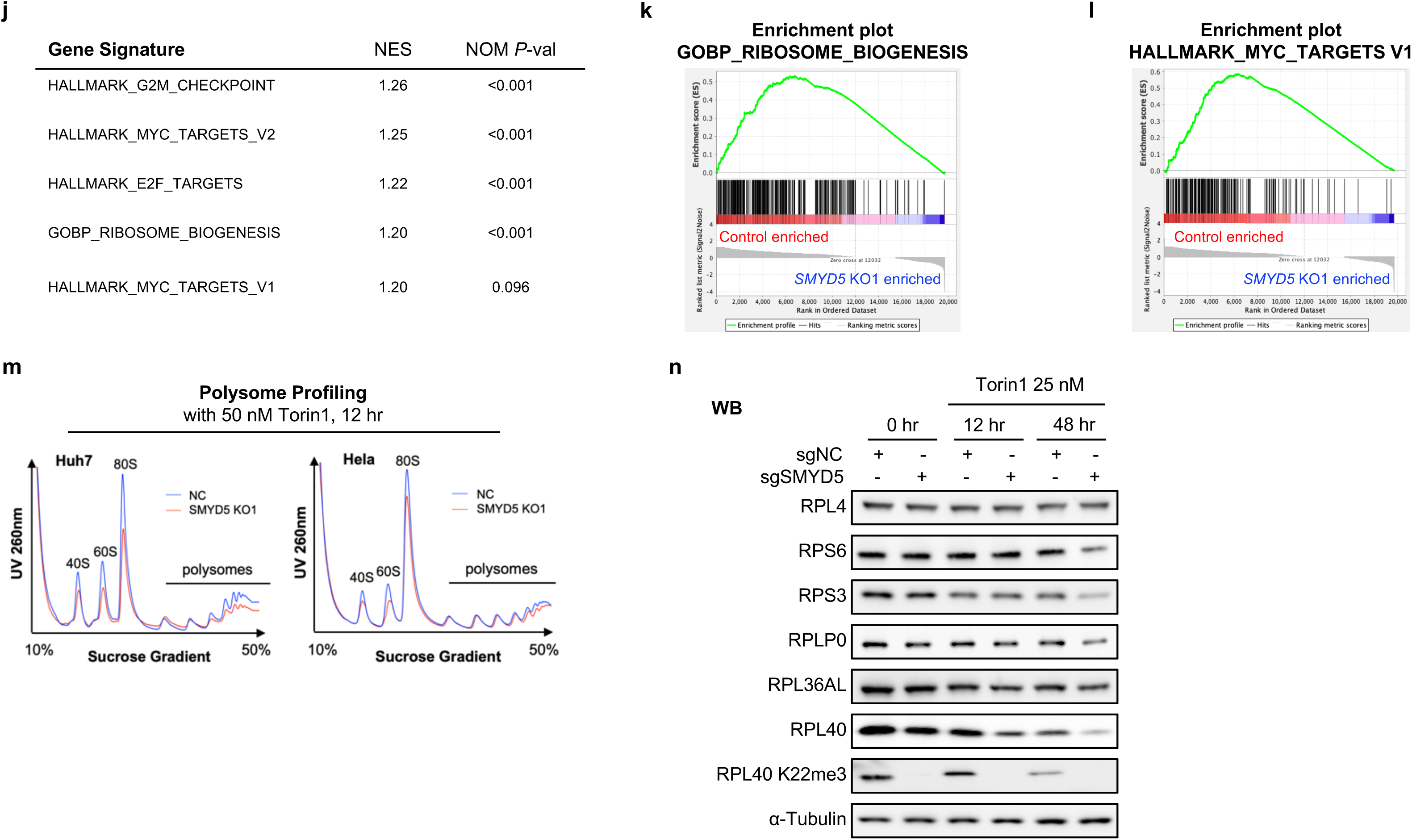
Related to Fig. 5. **a** Proliferation analyses of the control and SMYD5 KO of HeLa, Huh7, SNU449 and HepG2 cells. All data are represented as mean ± SD from three biological replicates. Two-tailed unpaired t-test. **b** Dose-response curves and IC50 measurement of the control and SMYD5 KO1 Huh7 cells treated with Torin1 (left) and Rapamycin (right) for 4 days. Data were represented as mean ± SD, and normalized to the untreated cells. Experiments were performed in duplication. **c** Proliferation analyses of the siNC and siRPL40 Huh7, with the indicated rescuing constructs and under the indicated situation (left and middle). The effect of rescue was verified by WB analyses (right). The proliferation analyses were performed in biological duplication as mean ± SD, *p < 0.05, ** p < 0.01, two-tailed unpaired T-test. **d** Proliferation analyses of the control and SMYD5 depleted (KO1) Huh7 and HeLa cells under the treatment of Rapamycin at the indicated concentrations. Experiments were performed three times. **e** WB analyses of newly synthesized proteins in the NC and *SMYD5* KO1 Huh7 cell lines with or without Torin1 treatment by AHA-click labeling labeling approach. **f-g** WB analyses of newly synthesized proteins in the NC and *SMYD5* KO1 SNU449 cell lines with or without Torin1 treatment by AHA-click labeling (**e**) and puromycin (**f**) labeling approach. **h** Scatterplots of ribo-seq showing translation efficiency (TE) correlation between NC and *SMYD5* KO1 Huh7 cells under Torin1 treatment. Genes with fold change > 1.5 or <1/1.5 were colored. **i** Codon bias analysis of ribo-seq between control and *SMYD5* KO1 Huh7 cells under Torin1 treatment. The fold changes of each codon occupancy were calculated, and the dashed lines marked 1.5-fold threshold. **j-l** Gene Set Enrichment Analysis (GSEA) identified enrichment of indicated gene sets in ribo-seq. Normalized enrichment scores (NES) and nominal *P* values (NOM *P*-val) were provided. **m** Polysome profilings of the control and SMYD5 depleted (KO1) Huh7 and HeLa cells under 50 nM Torin1 treatment for 12 hours. **n** WB analyses for some ribosome proteins abundance changes during Torin1 treatment in the NC and *SMYD5* KO1 Huh7 cell lines.

**Supplementary information, Fig. S7.**
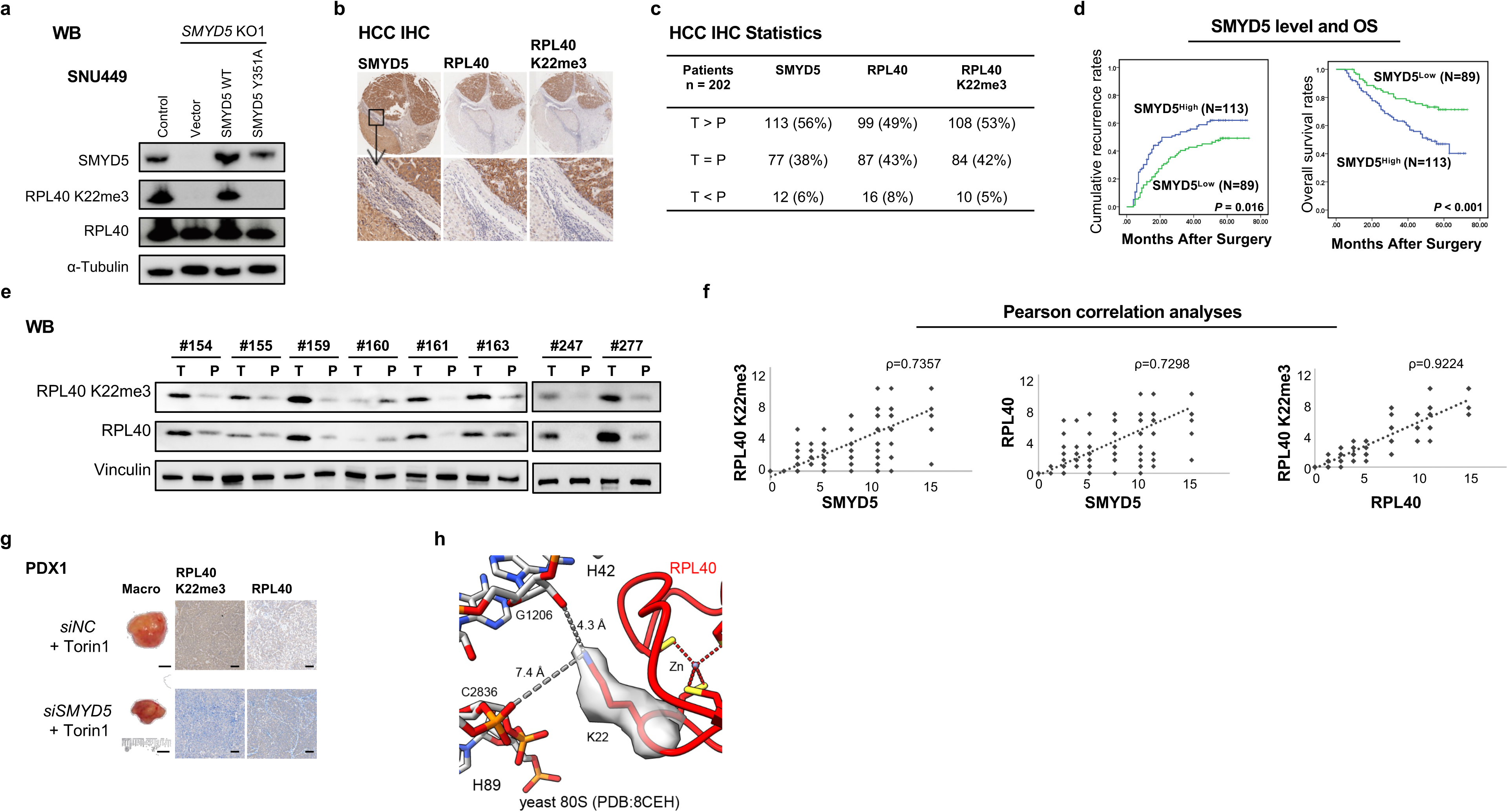
Related to Fig. 6 and discussion. **a** WB analyses of the indicated SMYD5, RPL40 and RPL40 K22me3 levels in the control, SMYD5 depletion and rescued SNU449 cells used for xenograft analyses in Fig. 6c. **b** IHC analyses of SMYD5, HIF1A, RPL40 and RPL40 k22me3 levels in human HCC tumor (T) samples and matched paratumor (P) tissues. Representative IHC views of 4 consecutive slides stained with indicated antibodies were shown in the upper panels; the corresponding enlarged closer views were shown in the lower panels. **c** The comparison of the IHC intensities of SMYD5, RPL40 and RPL40 K22me3 in 202 human HCC samples. **d** Kaplan-Meier’s analyses of the recurrency (left) and overall survival (right) of the 202 HCC patients divided by SMYD5 high and low (IHC intensities). **e** WB analyses of RPL40, RPL40 K22me3 and Vinculin (as control) levels in select HCC cases. T stands for tumor sample, and P stands for paired paratumor tissues. **f** Pearson correlation analyses of the IHC intensities. Left, the correlation of SMYD5 vs RPL40 K22me3; Middle, SMYD5 vs RPL40; Right, RPL40 vs RPL40 K22me3. The correlation coefficients (ρ) were indicated. *P < 2.2e-16*. **g** Representative gross images of PDX and immunohistochemical staining with the indicated antibodies of tumors from the PDX1 *siNC* + Torin1 group and *siSMYD5* + Torin1 group. Scale bars: 5 mm (whole mount) and 100 µm (histology). **h** RPL40 K22 residue is not methylated in yeast 80S ribosome (PDB: 8CEH). Residue K22 is shown in stick with its corresponding density map.

**Supplementary information, Fig. S8.**
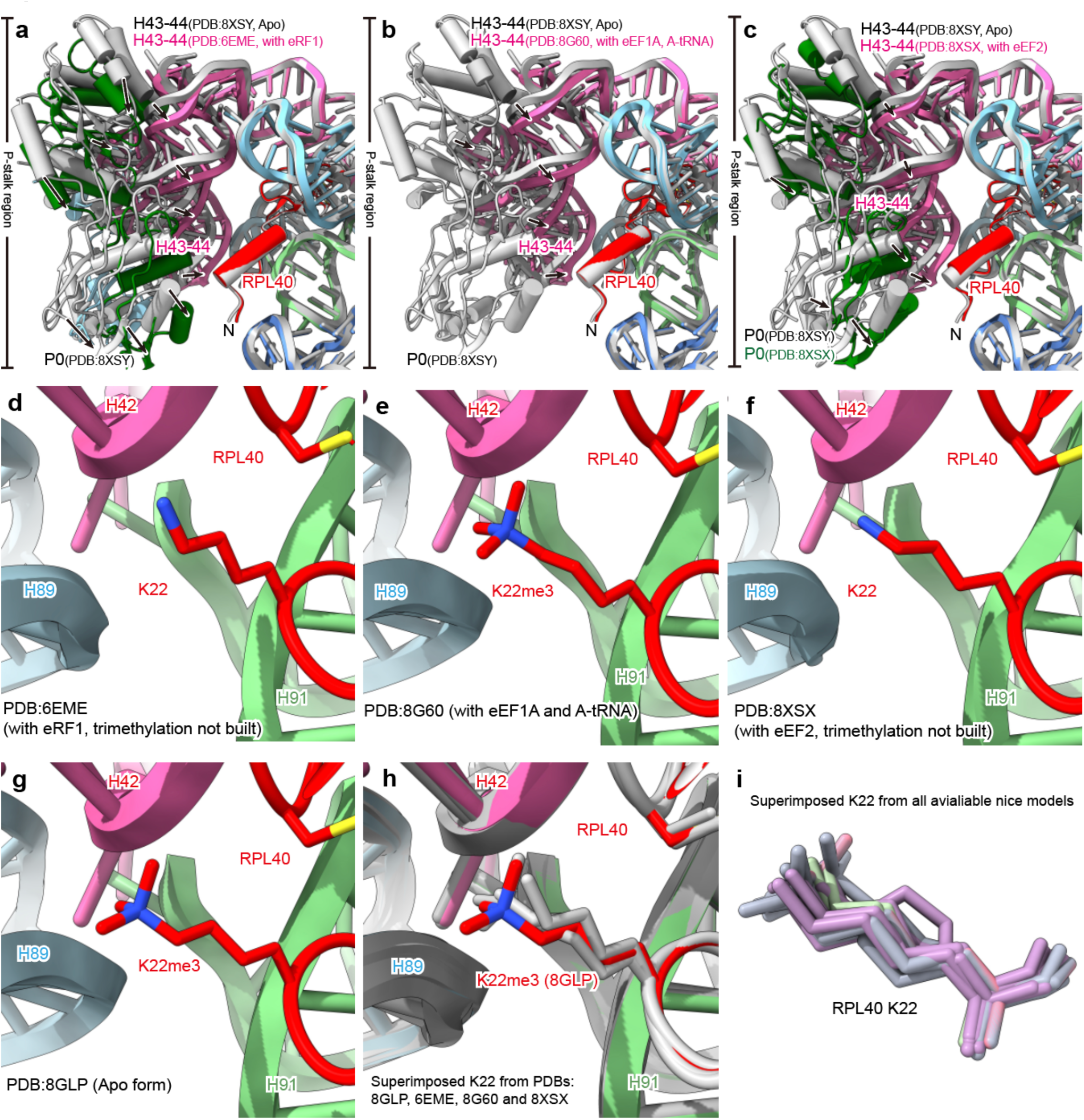
Structural comparisons among Apo ribosome and ribosomes complexed with translation factors eEF1A, eEF2, and eRF1 a-c. Overviews of RPL40 in the indicated ribosome structures. **d-h** Enlarged views of the RPL40 K22 side chains in the respective ribosome structures. **i** Across all comparisons, the positions of RPL40 and its K22 side chain exhibit no significant alterations, with variations attributable only to differences in model building. PDB numbers are provided for reference, and P stalk helices are highlighted in different colors.

## Notes

### Competing Interest Statement

The authors have declared no competing interest.

