## Supplementary material for "SMYD5 is a ribosomal methyltransferase that catalyzes RPL40 lysine methylation to enhance translation output and promote hepatocellular carcinoma": Materials and Methods: 4-Materials and Methods Final.docx

**Cell Culture, antibodies and general reagents**

HeLa, HEK293T, Huh7, HepG2, and SNU449 cells were cultured in Dulbecco’s Modified Eagle’s Medium (DMEM, Hyclone) supplemented with 10% fetal bovine serum (FBS, Gibco). HeLa, HEK293T, Huh7, and HepG2 cell lines were obtained from National Collection of Authenticated Cell Cultures (China), while SNU449 was obtained from ATCC.

For the RPL40 knockdown experiments, speciﬁc siRNA was synthesized (5’- CCUGCGAGGUGGCAUUAUU-3’) and introduced into the cells using RNAiMAX (Invitrogen) at approximately 50% conﬂuency. For the rescue experiment in Fig. S6c, the RPL40 coding sequence of CGCCTGCGAGGTGGCATTATT (wild type) was modified to CGGCTCCGGGGAGGGATCATC (seven synonymous mutations) to create RNAi resistance. Cells were collected at 60-96 hours post siRNA transfection for further analyses. For rescue experiments, pLenti-EF1a-BSD vector (Addgene) system was utilized. Lentiviruses were made in 293T cells and the viral supernatants were collected at 60 hour post-transfection and passed through a 0.45 μm filter prior to infection. After infection, HeLa and Huh7 cells were selected under 2 μg/mL Puromycin (Gibco) or 4 μg/mL Blasticidin (Gibco) for 5 days before cell proliferation analyses. For activating the phosphorylation of p38 and inhibiting ZAK in cells, anisomycin (ANS), homoharringtonine (HHT), harringtonine (HT), menadione(Mena) and M443 were obtained from MCE (HY-18982, HY-N0862, HY-14944, HY-B0332 and HY-112274).

Primary antibodies used in this study are listed in Supplementary Table S2.

**Generation of SMYD5 knockout cell lines and RPL40 K22R KI cell lines.**

CRISPR-Cas9 targeting system was utilized as previously described^1^. Guide RNA sequences for SMYD5 knockout are: KO1: 5’-CTGAGCAATACCACCAGGTC-3’, and KO2: 5’- AGCGCGGGTCTCCGTGGAAG-3’. Both sequences were designed to target exon1. Guide RNA sequence for RPL40 knock-in is: 5’-CATCGGAGCACACATACTTG-3’ and the donor template is:

5'-GCCAGCTGGCCCAGAAATACAACTGCGACAAGATGATCTGTCGCAGGTATGTGT

GCTCCGATGCTTGGGGGGCTGTGGGGGCTGCC-3'

**Cell fractionation separation**

Cells were swelled in hypotonic buffer (10 mM HEPES, pH 7.5, 1.5 mM MgCl_2_, 10 mM KCl, 0.5 mM DTT, with Cocktail protease inhibitors (Roche) and 1 mM PMSF) and incubated on ice for 20 min, and then treated with 0.2 % Triton X-100 for 5 min for lysis. Nuclei were collected by centrifugation at 2,000 rpm for 10 minutes at 4℃, and the supernatants were saved as cytoplasmic fraction.

**Immunofluorescence (IF) and immunoprecipitation (IP)**

Cells were seeded in 24-well plates at a density of 5 x 10^4^ cells/well at 24 hours before IF examination. Cytosol extracts used for IP and co-IP were prepared from HeLa cells and the experiments were carried out as described previously^2^.

**GST Pull-down assay**

Coding sequences of *SMYD5* and *RPL40* were cloned into pGEX4T-1 and GST tagged recombinant proteins were purified using affinity resins from SMART Lifesciences (Changzhou, China). Flag tagged SMYD5 and other recombinant methyltransferases were purchased from Active Motif China Inc. (Shanghai).

A total of 2.5 μg Flag-SMYD5 and 1 μg GST-RPL40 recombinant proteins were mixed and incubated with 150 μl binding buffer (20 mM TrisCl 7.4, 150 mM NaCl, and 0.2 % Triton X-100). Protein complex of GST-RPL40 and Flag-SMYD5 was immobilized by 10 ul GST resin and Flag-SMYD5 alone was set up as control. The resin and protein complexes were then washed with binding buffer for 4 times and subsequently subjected to SDS-PAGE examination and coomassie blue staining.

**Methyltransferase assay (MTase assay)**

Recombinant GST-SMYD5, Flag-SMYD5 and other indicated methyltransferases (Fig. 1a) were used as candidate enzymes. Recombinant GST-RPL40, RPL40 derived peptides (12-32 aa), histone tail derived peptides, mononucleosomes and the indicated cytosolic protein lysates were used as substrates. In general, 10 nM to 200 nM of individual methyltransferases were used and 5-10 μM substrates were used. For Fig. 2c and 2d, 20 μg of cytosolic protein lysates were incubated with 100 nM recombinant SMYD5. Methyltransferase buffer contained 20 mM TrisCl 8.0, 0.02% Triton X-100 and 0.5 mM TCEP with 10 μM SAM (or ^3^H-SAM) and assay were generally performed in a 10 - 20 μl reaction system, for 2 hours unless indicated.

The methyltransferase activities were monitored by four approaches, i.e. MALDI (Fig. 1b-c, 1f and Supplementary information, Fig. S2b,c), MS/MS after propionylation as previously described^3^ (Supplementary information, Fig. S2a), SDS-PAGE separation followed by radioautography (Fig. 1d-e, Fig. 2c-d and Supplementary information, Fig. S3f), and MTase-Glo Kit (Promega, Fig. 1a and Supplementary information, Fig. S2d). Kcat and Km analyses were performed by fitting Michaelis-Meanten equation with Graphpad. RPL40 peptides were synthesized from Qiangyao Biotech Co.,Ltd (Shanghai), and histone peptides were purchased from Active Motif China Inc. (Shanghai).

**Sample preparation for targeted mass spectrometry and PRM data acquisition**

To establish and optimize the PRM (Parallel Reaction Monitoring) method, we methylated the RPL40 (1-35) peptide in vitro by recombinant SMYD5. The methylation process was validated by MALDI-TOF. The unmethylated peptide segment was protected through propionylation and the peptide underwent reduction alkylation and trypsin digestion. Total protein samples from cells and tissues were lysed in SDS loading for SDS-PAGE gel electrophoresis. Approximately 100 μg total proteins of each sample were separated by electrophoresis. The RPL40 containing fractions were carefully excised from the gel at molecular weight around 10 kDa. The gel was then propionylated to protect unmodified residues, followed by in-gel reduction alkylation using 40 mM CAA (chloroacetamide) and 10 mM TCEP (tris (2-carboxyethyl) phosphine). In-gel digestion was performed using Trypsin (2 ng/μl) for 6-8 hours. Digested peptides were extracted using a solution of 50% acetonitrile and 0.1% trifluoroacetic acid (TFA). Then peptides were desalted using a C18 column and subsequently freeze-dried. Sample peptides were analyzed using on-line nanospray LC-MS/MS on an UltiMate 3000 system (Thermo Fisher Scientific, MA, USA) coupled to a timsTOF Pro mass spectrometer (Bruker Daltonics).

**MS analysis**

The data analysis was performed using SpectroDive 11.10 with default parameters. The software automatically corrected retention times and mass windows, and determined the optimal extraction window automatically. Peptide identification was conducted with a confidence threshold of Q value ≤ 0.01.

**Cell proliferation and chemical library screen**

For growth analyses of HeLa, Huh7 and HepG2 cells, a total of 1 x 10^5^ cells per well were seeded into 12 wells plate, the viabilities were measured at the indicated time point by CellTiter-Lumi^TM^ kit (Beyotime Inc). Cell viability was measured every 2 or 3 days and the cells were split to around 20 % density per well to ensure continuous growth at logarithmic phase. For chemical library screen, a total of 1,500 control and SMYD5 KO1 Huh7 cells per well were seeded in 96 well plates, cells with compound treatment were cultured for 4 days, and then subjected to the measurement of enhanced Cell-Counting Kit-8 activity (Beyotime Inc). Detailed information of individual compounds and concentrations used in the screen can be found in Supplementary information Table S2.

**Rescue experiment**

For RPL40 rescue experiment in Huh7 cells, we used siRNA to knockdown endogenous RPL40, and restored RPL40 expression by transiently transfecting siRNA resistant version of WT and K22R constructs simultaneously. Growth analyses were initiated 48 hours after the first transfection (set as Day 0). The second transfection was conducted in another 48 hours (Day 2 in the growth analyses) to sustain the knockdown of endogenous RPL40 and the ectopic expression of WT and K22R versions of RPL40.

**Soft Agar Assay**

A total of 500 cells per well were diluted in the top layer containing 2.5 % low viscosity methyl cellulose with complete medium (Life Technologies), and cultured on a bottom layer of 1% agar with complete medium in 12-well dishes. Both top and bottom layer medium were supplemented with 1% Penicillin-Streptomycin (Gibco) and 10% FBS. Additional media were added every 7 days to keep humidity before counting at day 25. The tumor volumes were collected and analysed by Image J (version = 1.53).

**Xenograft Assay**

For Huh7 xenograft assay, four-week-old female athymic nu/nu mice (BALB/c) housed under specific pathogen-free conditions were used in this study. A total of 2 x 10^6^ Huh7 (double check) cells were injected into the mammary fat pads of mice at the density of 2 x 10^6^ cells/ml. Mice were sacrificed and tumors were excised at day 18 after injection. Tumor development was measured every two days. Torin1 treatment and data collection were conducted as below. The animal care and experimental protocols were carried out in accordance with procedures and guidelines established by Shanghai Medical Experimental Animal Care Commission, all animal experiments were approved by Fudan University Institutional Committee.

For SNC449 xenograft assay, cells transduced with lentivirus expressing sgRNA/Cas9 and indicated reconstitution vectors expressing wildtype or mutant SMYD5 were transduced to immunocompromised 8-week-old NSG mice (NOD.SCID-IL2Rg-/-). The transplantation was performed by subcutaneous injection of cells mixed with matrigel (1:1) 2 ×10^6^ cells to the flanks of mice. When tumors became palpable, they were calipered every four days to monitor growth kinetics. Mice were treated as indicated with Torin1 (25 mg/kg once per day, IP) in vehicle 40% (2-hydroxypropyl)-β-cyclodextrin. Control animals underwent the same procedure but received vehicle treatment. All animals were numbered, and experiments were conducted in a blinded fashion. After data collection, treatment groups were revealed, and animals assigned to groups for analysis. Tumor size was measured using a digital caliper and tumor volume was calculated using the formula: Volume = (*width*)^2^ × *length* / 2 where *length* represents the largest tumor diameter and *width* represents the perpendicular tumor diameter. The endpoint was defined as the time at which a progressively growing tumor reached 20 mm in its longest dimension as approved by the MDACC IACUC protocol (00001636, PI: Mazur) and in no experiments was this limit exceeded.

**Patient-Derived Xenograft (PDX) Assay and siRNA delivery *in vivo***

PDXs were obtained from Liver Cancer Institute, Zhongshan Hospital, Fudan University, Shanghai, China and ethical approval was obtained from the research ethics committee of Fudan University affiliated Zhongshan Hospital, and written informed consent was obtained from each patient. Patient-derived xenograft models were established by transplanting small tumor fragments quickly and directly from surgical specimens of specific HCC patients into the flanks subcutaneous tissues of NSG mice. Excess tumor tissue can be frozen for the next inoculation. To knockdown the level of SMYD5 and RPL40 K22me3 in PDX tumors in situ, we delivered siRNA modified by cholesterol, phosphorothioate (PS), 2′-O-methyl (2′-OMe) and 2′-deoxy-2′-fluoro (2′-F) into tumors by intratumoral injection per three days from the beginning of measurement. Measurement were performed every two days when tumors were macroscopic. Tumor volume was calculated using the formula: Volume = (*width*)^2^ × *length* / 2 where length represents the largest tumor diameter and width represents the perpendicular tumor diameter.

**HCC animal model and IHC**

Reporter-tagged insertion with conditional potential *Smyd5^tm1a(EUCOMM)^* mouse strain was obtained from the European Mouse Mutant Archive repository^4^. Founder mice were crossed with *Rosa26*^FlpO^ deleter strain^5^ to generate conditional *Smyd5*^LoxP/LoxP^ allele.

*Alb^Cre^* mice have been described before^6^ and were obtained from the Jackson Laboratory (strain #003574). Mice were maintained on a mixed C57BL/6;129S1 strain background and we systematically used littermates as controls in all the experiments. For liver-specific deletion of SMYD5 we interbreed *Alb^Cre^* and *Smyd5^LoxP/LoxP^* mice. To establish HCC mouse model, a single dose of 1 mg/kg DEN (N-Nitrosodiethylamine, Sigma-Aldrich) was intraperitoneally injected (IP) into male mice at 2 weeks of age, followed by repeated administration of a low dose of a pro-fibrogenic agent carbon tetrachloride (CCl_4_, Sigma-Aldrich) at 0.2 ml/kg IP two times per week starting from 8 weeks of age for 12 weeks^7^. Six weeks after the last injection, mice were sacrificed for macroscopic and histopathological liver examination. All animals were numbered, and experiments were conducted in a blinded fashion. After data collection, genotypes were revealed, and animals assigned to groups for analysis. All mice were co-housed with littermates (2–5 per cage) in pathogen-free facility with standard controlled temperature of 22℃, with a humidity of 30–70%, and a light cycle of 12 h on/12 h off set from 7am to 7pm and with unrestricted access to standard food and water under the supervision of veterinarians, in an AALAC-accredited animal facility at the University of Texas M.D. Anderson Cancer Center (MDACC). Mouse handling and care followed the NIH Guide for Care and Use of Laboratory Animals. All animal procedures followed the guidelines of and were approved by the MDACC Institutional Animal Care and Use Committee (IACUC protocol 00001636, PI: Mazur).

Tissue specimens were fixed in 4% buffered formalin for 24 hours and stored in 70% ethanol until paraffin embedding. 3-μm sections were stained with hematoxylin and eosin (HE) or used for immunostaining studies. Immunohistochemistry (IHC) was performed on formalin-fixed, paraffin-embedded tissue (FFPE) sections using a biotin-avidin HRP conjugate method (Vectastain ABC kit). After incubated with 1^st^ antibodies, sections were developed with DAB and counterstained with hematoxylin. Pictures were taken using a PreciPoint M8 microscope equipped with the PointView software.

**Polysome profiling**

Polysome profiling was prepared as previously described^8^ with modifications. Approximately 2 x 10^7^ Huh7 and HeLa cells were incubated with 400 μM Cycloheximide (CHX, MCE) for 10 minutes, and then pelleted. Pellets were washed twice in PBS with 400 μM CHX and immediately lysed in 200 µl cold lysis buffer (100 mM KCl, 10 mM MgCl_2_, 50 mM TrisCl, pH 7.4 and 0.5 % NP-40) for 10 min on ice and pipetted to homogenize. The lysates were clarified by centrifugation at 700g for 5 min at 4 °C to discard cell nucleus and 12,000 g for 10 min at 4 °C to discard mitochondria and debris. Lysates were then loaded onto 10–50 % sucrose gradients and ultracentrifuged in a SW41 Ti swinging-bucket rotor (331362, Beckman) at 36,000 rpm for 2h at 4 °C. For RNase A digested polysome profiling^9^, lysates containing 100μg of total RNA were treated with RNase A (Thermo Fisher Scientific) at 5 mg/L for 45 min at RT and added 400U of RNase Inhibitor (Beyotime Inc) to terminated the digest reaction. Digested lysates were then loaded onto 10–40 % sucrose gradients and ultracentrifuged in a SW41 Ti swinging-bucket rotor (331362, Beckman) at 36,000 rpm for 2h at 4 °C. Samples were fractionated using a Biocomp gradient fractionator for absorbance polysome profiles and separation. Control and SMYD5 KO samples were measured and loaded evenly with equivalent 260 nM OD values.

**Ribosome profiling sequencing (Ribo-seq) and data processing**

Ribo-seq was performed as previous described^10^ and ARTseq Ribosome Profiling Kit’s instructions with modifications. About 10^7^ cells were pre-treated by 100 μg/ml cycloheximide (MCE) for 10 min at 37℃, then washed and collected by ice-cold PBS containing 100 μg/ml cycloheximide. Cells were lysed by 120 μl Mammalian Polysome Buffer (10 mM Tris-HCl pH 7.5, 100 mM KCl, 5 mM MgCl2, 1% Trion X-100 with 1x protease inhibitor cocktail (Roche)) for 10 min. After centrifugation at 13,000 g for 10min at 4℃. 10 ul lysate was kept for mRNA-seq and purified by RNA Clean & Concentrator-5 kit (RCC-5 R1016, zymo), followed by rRNA removal with Ribo-off rRNA Depletion (N406, Vazyme) and library preparation with VAHTS Universal RNA-seq Library Prep Kit (NR605, Vazyme). Add about 30U RNase I (EN0601, Thermo) to 100ul lysate and digest for 45 min at room temperature. Digestion was stopped by the addition of 4 μl of Superase-In (AM2696, Thermo). Meanwhile, MicroSpin S-400 HR columns (27514001, Cytiva) were equilibrated with 3 ml of Mammalian Polysome Buffer by gravity flow and emptied by centrifugation at 600g for 4 min. We then immediately loaded 100 μl of the digested lysate on the column and eluted the column by centrifugation at 600g for 2 min. RNA was extracted by RCC-5 and separated on 15% denaturing urea-PAGE gel. After SYBR gold (S11494, Thermo) staining, the size ranges from 25 nt to 40 nt was cut out and recovered by small-RNA PAGE Recovery Kit (R1070, zymo). The eluted RNA were mix with Superase-In, T4 PNK and Buf A (EK0031, Thermo) at 37 ℃ for 15 min and supplemented by 1 mM ATP (R0441, Thermo) for another 30 min before extraction by RCC-5 and generation library by Small RNA Library Prep Kit (NR811, Vazyme). The libraries were sequenced by NovaSeq 6000 (conducted by Nanjing Gaoxin Precision Medicine Technology Co., Ltd).

After sequencing, reads were trimmed by TrimGalore v0.6.10 to excise low-quality bases and adaptors in both mRNA and Ribo-seq datasets. Quality assessment was subsequently conducted utilizing FastQC. Contamination originating from ribosomal RNA was discerned through alignment of reads to human rRNA sequences employing Bowtie2^11^ v2.3.5.1, followed by the exclusion of mapped reads from subsequent analyses.The human genome reference sequence (GRCh38.p14.genome.fa) and annotation files (gencode.v45.chr_patch_hapl_scaff.annotation.gtf) were acquired from the GENCODE browser. Within the annotation file, the longest isoforms were retained using AGAT v1.2.1 (agat_sp_keep_longest_isoform.pl), and only the canonical chromosomes (1-22, X, Y) were selected in the sequence file. The remaining Ribo-seq reads were aligned to the filtered human reference genome using STAR^12^ v2.7.10b with specified parameters (--runMode alignReads --readFilesCommand zcat --quantMode TranscriptomeSAM GeneCounts --twopassMode Basic). RNA-seq clean reads were analyzed using a combination of HISAT2^13^ v2.2.1 and StringTie^14^ v2.1.7. Raw mRNA counts were normalized using DESeq2^15^, with exclusive retention of highly expressed genes (Average FPKM in NC and KO > 2.5). Ribosome Protected Footprint (RPF) reads were normalized relative to the cumulative counts of mitochondrial RPFs^16^. Translation efficiency (TE) was determined by dividing RPF counts by mRNA counts. Statistical analysis of differentially transcribed genes within each sample pair was executed using the R package Xtail^17^. Codon bias for each group was computed utilizing CONCUR^18^ v1.0, while codon occupancy was quantified and depicted through customized Python scripts. Read density pertaining to focused transcripts was calculated employing RiboMiner^19^ v0.2 and visualized using custom Python scripts. GSEA analysis were performed by GSEA software v4.3.3 based on hallmark gene sets and ribosome biogenesis gene set^20,21^.

**HCC tissue microarray and IHC**

A total of 202 formalin-fixed paraffin embedded HCC tissues (containing tumor and paratumor compartments) were collected from consecutive patients with HCC who underwent curative resection from 2006 to 2007 at the Liver Cancer Institute of Fudan University (Shanghai, China). Histopathological diagnoses were based on World Health Organization criteria. Ethical approval was obtained from the research ethics committee of Fudan University affiliated Zhongshan Hospital, and written informed consent was obtained from each patient. Among the 202 patients,172 cases were male, 30 cases were female. The age distribution was from 27 to 84 years old. Tumor size ranged from 0.5 cm to 20 cm. Follow-up data were summarized at the end of December 2013, with a median follow-up of 51 months (range: 5-73 months). Tissue microarrays (TMAs) were constructed and immunohistochemistry (IHC) was performed as previously described elsewhere^22^.

**Statistical Analysis**

For cell proliferation assays, all statistical data were calculated using GraphPad Prism 9. Comparisons of data were performed by two-sided, parametric and unpaired t-test; P values of less than 0.05 were considered significant. n=3 or 4 (the number of samples) for each experimental group. Values are presented as mean ± standard deviation (SD). 3 biological replicates were performed and one representative was shown.

For soft agar assays, the statistical data were calculated using GraphPad Prism 9. Comparisons of data were performed by two-sided, parametric and unpaired t-test; P values of less than 0.05 were considered significant. n=3 (the number of samples) for each experimental group. Values are presented as mean ± standard deviation (SD). 3 biological replicates were performed and one representative was shown.

For IHC experiments, statistical analysis was performed with SPSS software (19.0; SPSS, Inc., Chicago, IL). Values are presented as mean ± standard deviation (SD). The Student t test was used for comparisons between groups. Pearson’s correlation analyses were performed for the IHC intensities of SMYD5, RPL40, RPL40 K22me3 and HIF1A. Overall survival rates were analyzed using Kaplan-Meier’s method and the log-rank test. *P* < 0.05 was considered statistically signiﬁcant.

**Data availability**

High-throughput sequencing data were deposited to GSE241588.
