## Supplementary information Table S1 for "SMYD5 is a ribosomal methyltransferase that catalyzes RPL40 lysine methylation to enhance translation output and promote hepatocellular carcinoma": Supplementary information Table S1.pdf

**Supplementary information, Table S1. Antibodies used in this study.**

| Name | Source | Identifier | Dilutions |
|---|---|---|---|
| **WB** | | | |
| SMYD5 | ABclonal | Cat#A6191 | 1 : 1000 |
| SMYD5 | Sigma | Cat#HPA015514 | 1 : 500 |
| Lamin B1 | Proteintech | Cat#66095-1-lg | 1 : 5000 |
| α-Tubulin | Proteintech | Cat#66031-1-Ig | 1 : 5000 |
| RPL40 | Abcam | Cat#ab109227 | 1 : 10000 |
| RPL40 K22me3 | custom generated | | 1 : 2000 |
| Phospho-p38 | Proteintech | Cat#28796-1-AP | 1 : 10000 |
| RPL4 | Proteintech | Cat#67028-1-lg | 1 : 5000 |
| RPS3 | Proteintech | Cat#11990-1-AP | 1 : 5000 |
| RPS6 | Proteintech | Cat#14823-1-AP | 1 : 5000 |
| Vinculin | Cell Signaling Technology | Cat#13901 | 1 : 2000 |
| Flag | Sigma | Cat#F3165 | 1 : 5000 |
| Streptavidin -HRP | Life Technologies | Cat#434323 | 1 : 1000 |
| **IF** | | | |
| SMYD5 | ABclonal | Cat#A6191 | 1 : 200 |
| HA | Cell Signaling Technology | Cat#3724 | 1 : 200 |
| DAPI | solarbio | Cat#C0060 | 1 : 1000 |
| **IP** | | | |

| | | | |
|---|---|---|---|
| Anti-FLAG M2 affinity gel | Sigma | Cat#A2220 | |
| RPL40 | Abcam | Cat#ab109227 | 1 : 100 |
| SMYD5 | ABclonal | Cat#A6191 | 1 : 100 |
| Normal rabbit IgG | Santa Cruz Biotechnology | Cat#sc-2027 | |
| **IHC** | | | |
| SMYD5 | ABclonal | Cat#A6191 | 1 : 250 |
| SMYD5 | Sigma | Cat#HPA015514 | 1 : 100 |
| pH3 | Cell Signaling Technology | Cat#9701 | 1 : 1000 |
| RPL40 | Abcam | Cat#ab109227 | 1 : 500 |
| RPL40 K22me3 | custom generated | | 1 : 2000 |
