## Supplementary information Table S2 for "SMYD5 is a ribosomal methyltransferase that catalyzes RPL40 lysine methylation to enhance translation output and promote hepatocellular carcinoma": Supplementary information Table S2.pdf

**Supplementary information, Table S2. Compounds used in drug library screen.**

| Compound Name | Working Concentration | Inhibitory Ratio (*SMYD5* KO1/NC) |
|---|---|---|
| Olaparib (AZD2281, Ku-0059436) | 200nM | 0.911319504 |
| Odanacatib (MK-0822) | 200nM | 1.031316901 |
| Tanespimycin (17-AAG) | 200nM | 1.087523552 |
| Anastrozole | 200nM | 1.087893896 |
| Aprepitant | 200nM | 1.04457781 |
| Ispinesib (SB-715992) | 200nM | 0.901058343 |
| Zibotentan (ZD4054) | 100nM | 1.02918495 |
| GSK429286A | 100nM | 0.981534395 |
| Adavosertib （MK-1775） | 200nM | 0.910247007 |
| Ambrisentan | 200nM | 1.044900009 |
| Ibrutinib (PCI-32765) | 200nM | 0.9665821 |
| Turofexorate Isopropyl (XL335) | 200nM | 1.062983581 |
| Nepicastat (SYN-117) HCl | 200nM | 0.907145659 |
| Crenolanib (CP-868596) | 200nM | 1.024199874 |
| CHIR-98014 | 200nM | 1.080231542 |
| LDC1267 | 200nM | 0.96411533 |
| Torin 2 | 50nM | 1.035980404 |
| Carfilzomib (PR-171) | 50nM | 1.030724678 |

| | | |
|---|---|---|
| GNF-2 | 200nM | 0.993039962 |
| JNJ-7777120 | 200nM | 1.111280896 |
| Eprosartan Mesylate | 200nM | 1.013748036 |
| JNK-IN-8 | 200nM | 0.948640249 |
| Rolapitant | 200nM | 0.977884563 |
| GSK3787 | 200nM | 0.861638195 |
| Rasagiline | 200nM | 0.913740278 |
| JANEX-1 | 200nM | 1.006499707 |
| VX-702 | 200nM | 0.847257432 |
| BI-4464 | 200nM | 0.828744004 |
| PP2 | 200nM | 0.842773251 |
| GSK2656157 | 200nM | 0.94394401 |
| SGC 0946 | 200nM | 0.992140279 |
| GSK2334470 | 200nM | 0.89811131 |
| AGI-5198 | 200nM | 0.943777222 |
| Bisindolylmaleimide I (GF109203X) | 200nM | 0.931795451 |
| Alvelestat (AZD9668) | 200nM | 0.916020796 |
| UNC0642 | 200nM | 0.947224791 |
| AGI-6780 | 200nM | 0.986106111 |
| PFI-2 HCl | 200nM | 0.904432403 |
| Poziotinib (HM781-36B) | 200nM | 0.912321086 |

| | | |
|---|---|---|
| LDC000067 | 200nM | 0.901922672 |
| FTI 277 HCl | 200nM | 1.143453816 |
| Nexturastat A | 200nM | 0.938202191 |
| Santacruzamate A (CAY10683) | 200nM | 1.054061188 |
| Picropodophyllin (PPP) | 200nM | 1.23202818 |
| BPTES | 200nM | 1.062369516 |
| Emricasan | 200nM | 0.860269874 |
| EPZ020411 2HCl | 200nM | 1.055030269 |
| SBI-0206965 | 200nM | 0.97055621 |
| Venetoclax (ABT-199, GDC-0199) | 200nM | 1.141362159 |
| SB366791 | 200nM | 1.044741028 |
| Etomoxir (Na salt) | 200nM | 0.988544233 |
| Selonsertib (GS-4997) | 200nM | 0.91920362 |
| T-3775440 HCl | 200nM | 0.93390312 |
| BAY-876 | 200nM | 1.14585723 |
| GSK'872 (GSK2399872A) | 200nM | 0.984042747 |
| LY3214996 | 200nM | 1.051760245 |
| Z944 | 200nM | 1.103401039 |
| DMAT | 200nM | 1.15417303 |
| pm26TGF-β1 peptide (TFA) | 200nM | 1.12758108 |
| Bemcentinib (R428) | 200nM | 1.021975702 |

| | | |
|---|---|---|
| FF-10101 | 200nM | 1.005815614 |
| ETC-206 (AUM 001) | 200nM | 1.063205602 |
| GSK467 | 200nM | 1.058399859 |
| TH588 | 200nM | 0.969350794 |
| BX-795 | 200nM | 1.01857296 |
| Cediranib (AZD2171) | 200nM | 1.029825411 |
| SHP099 | 200nM | 0.990923371 |
| AZD7648 | 200nM | 1.057386037 |
| SMI-4a | 200nM | 1.174798759 |
| GSK461364 | 50nM | 1.153861447 |
| Pelitinib (EKB-569) | 50nM | 1.036961433 |
| RS504393 | 200nM | 1.019870705 |
| Omipalisib | 50nM | 0.655857915 |
| UK-371804 HCl | 200nM | 1.029559455 |
| AMG-458 | 200nM | 1.096898209 |
| Daidzin | 200nM | 1.072568194 |
| Finerenone | 200nM | 1.037885672 |
| TP-064 | 200nM | 0.970002315 |
| AM966 | 200nM | 1.009542205 |
| IRAK4-IN-1 | 200nM | 1.237955859 |
| IOX2 | 200nM | 1.108777258 |

| | | |
|---|---|---|
| Canagliflozin | 200nM | 1.08162081 |
| Palbociclib (PD-0332991) HCl | 200nM | 0.967082988 |
| bpV (HOpic) | 200nM | 1.043614958 |
| VTP50469 | 200nM | 1.0506594 |
| WQ 1 | 200nM | 1.044233596 |
| Pitolisant hydrochloride | 200nM | 1.078089194 |
| Linagliptin | 200nM | 0.934215605 |
| ZT-12-037-01 | 200nM | 0.857211151 |
| HS-276 | 200nM | 0.933047447 |
| NVP-CGM097 | 200nM | 0.951346876 |
| HG-9-91-01 | 200nM | 1.033892037 |
| ML347 | 200nM | 0.952825987 |
| Alvimopan | 200nM | 1.067622754 |
| Diphenyleneiodonium chloride (DPI) | 200nM | 0.811540995 |
| GSK'963 | 200nM | 0.934410954 |
| GSK'547 | 200nM | 1.051551463 |
| SSR128129E | 200nM | 0.977970886 |
| TED-347 | 200nM | 1.016815573 |
| MLi-2 | 200nM | 1.228831438 |
| T-26c | 200nM | 1.114705998 |
| Larotrectinib | 200nM | 1.114025767 |

| | | |
|---|---|---|
| PTC-209 HBr | 200nM | 0.985129147 |
| CITCO | 200nM | 1.220365609 |
| Tegoprazan | 200nM | 1.060317393 |
| RO 5028442 (RG7713) | 200nM | 1.039736072 |
| SC 236 | 200nM | 1.013045427 |
| A-803467 | 200nM | 1.039916991 |
| Torin1 | 100nM | 0.551457552 |
| Trametinib | 100nM | 0.840932661 |
| Rapamycin | 500nM | 0.726116714 |
| Alpelisib | 50nM | 0.826085523 |
| Gefitinib | 50nM | 0.848285017 |
| ADU-S100 | 50nM | 0.987128941 |
| Cenicriviroc | 50nM | 0.927603116 |
| 3-Bromopyruvic acid | 50nM | 0.914605572 |
| Galunisertib | 50nM | 1.103435781 |
| Ipatasertib | 50nM | 0.933878351 |
| Crizotinib | 50nM | 0.942497601 |
| WZB117 | 50nM | 0.942497601 |
| Z-DEVD-FMK | 50nM | 1.040792957 |
| AZD-5069 | 50nM | 0.963056461 |
| GSK269962A | 50nM | 0.994255745 |

| | | |
|---|---|---|
| Stattic | 50nM | 0.937989569 |
| AZD-7762 | 50nM | 0.958145326 |
| FX-11 | 50nM | 1.028966413 |
| Osimertinib | 50nM | 1.067055123 |
| Dasatinib | 50nM | 0.965696766 |
| SB 203580 | 50nM | 0.904752869 |
| TBHQ | 50nM | 0.894304265 |
| Pirfenidone | 50nM | 0.915560856 |
| Vemurafenib | 50nM | 0.945527567 |
| C75 | 50nM | 0.941983355 |
| AT9283 | 50nM | 1.023221366 |
| Honokiol | 50nM | 0.971931513 |
| CHIR-99021 | 50nM | 1.017841474 |
| KIN1148 | 50nM | 0.925154839 |
| Ruxolitinib | 50nM | 0.957365982 |
| Wogonin | 50nM | 0.946882016 |
| Lonidamine | 50nM | 1.041984023 |
| Tozasertib | 50nM | 0.988323275 |
| ABT-737 | 50nM | 0.980623809 |
| Mitapivat | 50nM | 0.974695733 |
| Nifurtimox | 50nM | 0.925370008 |

| | | |
|---|---|---|
| Amlexanox | 50nM | 1.095645984 |
| Nutlin-3a | 50nM | 1.062771737 |
| Venetoclax | 50nM | 0.976717877 |
| Forsythoside B | 50nM | 0.945893162 |
| FH535 | 50nM | 0.987395881 |
| EMT inhibitor-1 | 50nM | 1.0152316 |
| Dorsomorphin | 50nM | 0.879552854 |
| Firsocostat | 50nM | 0.973075521 |
| Birinapant | 50nM | 0.983162037 |
| SCH772984 | 50nM | 1.085113111 |
| TEPP-46 | 50nM | 1.080685413 |
| BAY 11-7082 | 50nM | 0.936045092 |
| V-9302 (hydrochloride) | 50nM | 0.961269473 |
| Ripasudil | 50nM | 1.098428286 |
| Lorlatinib | 50nM | 0.965577588 |
| Asiaticoside | 50nM | 0.977396539 |
| Cinacalcet | 50nM | 0.936203677 |
| Epacadostat | 50nM | 1.017436959 |
| SB 202190 | 50nM | 1.056197833 |
| SKL2001 | 50nM | 0.883960965 |
| Fatostatin | 50nM | 0.836804421 |

| | | |
|---|---|---|
| PF-06928215 | 50nM | 1.058183257 |
| Staurosporine | 50nM | 0.579757716 |
| XMU-MP-1 | 50nM | 0.945015897 |
| A 1070722 | 50nM | 0.955976833 |
| Erlotinib | 50nM | 0.924302698 |
| Sorafenib | 50nM | 0.87954172 |
| Prexasertib | 50nM | 0.782503237 |
