## Supplementary information Table S3 for "SMYD5 is a ribosomal methyltransferase that catalyzes RPL40 lysine methylation to enhance translation output and promote hepatocellular carcinoma": Supplementary information Table S3.pdf

***Smyd5* knockout Mouse Phenotype Summary**

| **Mouse Phenotypes** | **Male** | **Female** |
|---|---|---|
| Mortality/aging | n.s. | n.s. |
| Growth/size/body region | n.s. | Increased lean body mass |
| Reproductive system | n.s. | n.s. |
| Homeostasis/metabolism | n.s. | n.s. |
| Cardiovascular system | n.s. | n.s. |
| Digestive/liver/biliary system | n.s. | n.s. |
| Behavior/neurological system | Abnormal Gait | Abnormal Gait |
| Renal/urinary system | n.s. | n.s. |
| Immune system | n.s. | n.s. |
| Limbs/digits/tail | n.s. | n.s. |
| Skeleton | n.s. | Abnormal bone mineralization |
| Integument or pigmentation | n.s. | n.s. |
| Craniofacial | n.s. | n.s. |
| Hearing/vestibular/ear | n.s. | n.s. |
| Endocrine/exocrine gland | n.s. | n.s. |
| Vision/eye | n.s. | n.s. |

Supplementary information, Table S3. Table Summary of *Smyd5* KO mice phenotype recorded on the website of **International Mouse Phenotyping Consortium**. URL: www.mousephenotype.org
